# Multi-dimensional optimization of a lysin towards a ribolysin against life-threatening *S. aureus* infections: Fc-LysM-CHAP and its strong synergy with standard of care antibiotics

**DOI:** 10.64898/2026.04.27.720530

**Authors:** Adriana Badarau, Benedikt Bauer, Zehra Visram, Renping Qiao, Andrea Majoros-Hashempour, Johannes Söllner, Svetlana Durica-Mitic, Orla M. Dunne, Robert Kluj, Donaat Kestemont, Ann-Katrin Kieninger, Monika Kulig, Rocío Berdaguer, Timo Schwebs, Manuel Zerbs, Philipp Czermak, Marina von Freyberg, Julia Schmidt, Laura Schmidberger, Dominik Mayer, Maria Protano, Hannah A. Miller, Gavin M. Palowitch, Charles L. Dulberger, Maria Massaro, Nikita Ivanisenko, Hippolyte Jacomet, Didier Ngatcha Bakoue, Francesco Oteri, Gali Sela, Lorenzo Corsini

**Author notes:** Corresponding authors: Lorenzo Corsini, Adriana Badarau (;). Contributed equally (Benedikt Bauer, Zehra Visram).

## Abstract

Bacterial lysins are promising novel antimicrobials but are limited by poor pharmacokinetics and challenging manufacturability. We developed a lysin discovery platform tailored for lysin delivery via mRNA: staphylococcal LysM-CHAP (cysteine, histidine-dependent amidohydrolases/peptidase) autolysin was selected and its serum half-life extended via Immunoglobulin G1-Fc-fusion. The Fc-induced drop in lysin potency was rescued by the concerted optimization of linkers, binding kinetics and catalytic activity, using a combination of rational and AI-guided approaches. The engineered Fc-LysM-CHAP was active against planktonic bacteria (minimum inhibitory concentration of 1 – 2 µg/mL) and simulated endocardial vegetations and synergized strongly (Fractional eradication concentration index FECI = 0.06) with cell wall active antibiotics in vitro. In mouse models of Staphylococcus aureus sepsis, the recombinant Fc-LysM-CHAP - antibiotic combination was superior to single agent treatments and mRNA-delivered Fc-LysM-CHAP showed single agent activity at a mRNA-lipid nanoparticle dose as low as 0.2 mg/kg.

## Main text

Staphylococcus aureus (Sa) causes a broad spectrum of clinical manifestations, from mild skin and soft tissue infections to life-threatening invasive disease including bacteraemia, endocarditis, osteomyelitis, pneumonia and device-related infections driven by biofilm formation. Sa ranks as one of the leading bacterial causes of death, responsible for ∼1 million deaths per year globally^1^.

The evasion of antibiotic action and immune surveillance stems from Sa adaptation to the human host: biofilm formation^2^, spreading to vital organs, causing tissue damage by releasing toxins^3^, blunting the adaptive immunity by binding immunoglobulins^4^ and surviving both extra and intra-cellularly^5^. Our aim was to develop direct lytic agents (bacterial lysins) as a new therapeutic modality, to counteract as many of these evasion mechanisms as possible.

Bacteriophage-derived endolysins and the related autolysins - cell wall hydrolases that cleave specific peptidoglycan bonds - have emerged as a novel class of antibacterial candidates fundamentally distinct from small-molecule drugs. Lysins display a modular architecture with one or more cell wall binding and catalytic domains connected by flexible linkers. They trigger rapid bacteriolysis independently of metabolic state, show low propensity for resistance development and exhibit potent activity against biofilm-embedded cells, where small molecule antibiotics are mostly silent^6,7^. Examples include exebacase (CHAP-SH3), an anti-staphylococcal lysin that has progressed to Phase 3 clinical trials for the treatment of staphylococcal bacteraemia and endocarditis, but was discontinued due to lack of efficacy^8,9^, and lysostaphin (PepM23-SH3)^10^, one of the most potent anti-staphylococcal lysins, and the first to be used in the clinic^11^.

However, native lysins are characterized by rapid renal clearance and short serum half-lives due to their small size (20-30 kDa, close to renal filtration limit) and positive charge. This necessitates frequent dosing or half-life extension strategies that may compromise clinical activity^12,13^.

Here we applied rational protein engineering to optimize the developability of LysM-CHAP, a natural autolysin, making it suitable for mRNA-based delivery and thus improving manufacturability. The engineered lysin candidate was encoded by a nucleoside-modified mRNA formulated in lipid nanoparticles (LNP) - a platform similar to the modRNA-LNP technology used for the COVID-19 vaccine, BNT162b2 (Comirnaty®)^14^. The lysin is highly potent, shows synergy with cell wall active antibiotics (vancomycin, cefazolin) in vitro, and displays good pharmacokinetics and efficacy in mouse models of Sa sepsis when delivered as recombinant protein or mRNA-LNP that, with different payloads and different LNP, has been tolerated in humans^15–17^.

## Results

### Half-life extension

A screen of ∼150 natural lysins with phage or bacterial origin led to the identification of the LysM-CHAP architecture as highly enriched among lysins active against Sa^18^. The C-terminal LysM-CHAP fragment of autolysin LytN (amino acids 158-372) was selected for optimization. LytN is involved in cell wall remodelling during Sa cell division^19,20^. It binds to repeating disaccharide β-N-acetylmuramic acid-(1→4)-β-N-acetylglucosamine (MurNAc-GlcNAc) peptidoglycan polymers via its LysM domain, recruiting the CHAP domain to its substrate. The CHAP domain then cleaves the peptidoglycan stem peptide, acting both as an amidase (on MurNAc-L-Ala) and peptidase (on D-Ala-Gly)^19^. The parental LytN fragment (L0) showed low thermal stability and degraded rapidly in human serum. We applied several rounds of directed evolution, by generating high diversity lysin libraries which were simultaneously displayed and secreted from the same yeast cell, followed by selection of clones for yeast cell surface expression and for activity of selected yeast colonies on agar plates containing autoclaved Sa cells. We further subjected the main lineages from round 1 and 2 (L1 and L2, respectively) to rational design, yielding a set of variants with up to 18 amino-acid substitutions vs L0 (Extended Data Table 1). The variants displayed significantly decreased charge, increased thermal stability, good levels of secretion from human embryonic kidney (HEK) cells, and were stable in human serum, while maintaining activity against Sa (minimal inhibitory concentration [MICs] of 0.25-2 µg/mL vs 0.5 µg/mL for L0) (Extended Data Table 1). To increase the half-life of LysM-CHAP, we selected L1, which outperformed L2 in terms of stability and secretion (Extended Data Table 1), and fused it to human IgG1-Fc using a set of different linkers, yielding a library of Fc-LysM-CHAP variants, followed by screening in HEK supernatants for secretion and activity. The linker sequence had a substantial impact on expression levels and homogeneity of secreted proteins, with e.g. glycine-rich linkers associated with no or little full-length protein in supernatants. The helical “hel8”-linker (A-(EAAAK)_4_-ALEA-(EAAAK)_4_-A) displayed good secretion and bactericidal activity in the absence of any visible free lysin release (Extended Data Table 2). Similar screens for L1 fusions to other pharmacokinetic (PK)-enhancing moieties using hel8 (Fig. 1a) failed to identify active constructs (e.g. human serum albumin (HSA)^21^, binders to HSA^22^ or to IgG1-Fc)^23^ or did not translate to an improved PK in vivo. C-terminal Fc fusions (LysM-CHAP-Fc) were also inactive (data not shown). Fc-hel8 fusions with additional LysM-CHAP variants were designed and purified from HEK supernatants (Extended Data Fig. 1a, b). Consistent with the supernatant screen, growth inhibition assays showed ∼16-32-fold increased MICs (decreased potency), compared to the naked LysM-CHAP counterparts (Fig. 1b); there was a trend in net positive charge improving potency (i.e. decreasing MIC; significant Spearman correlation, r=-0.8182, P=0.0169) (Extended Data Fig. 1c)

**Figure 1:**
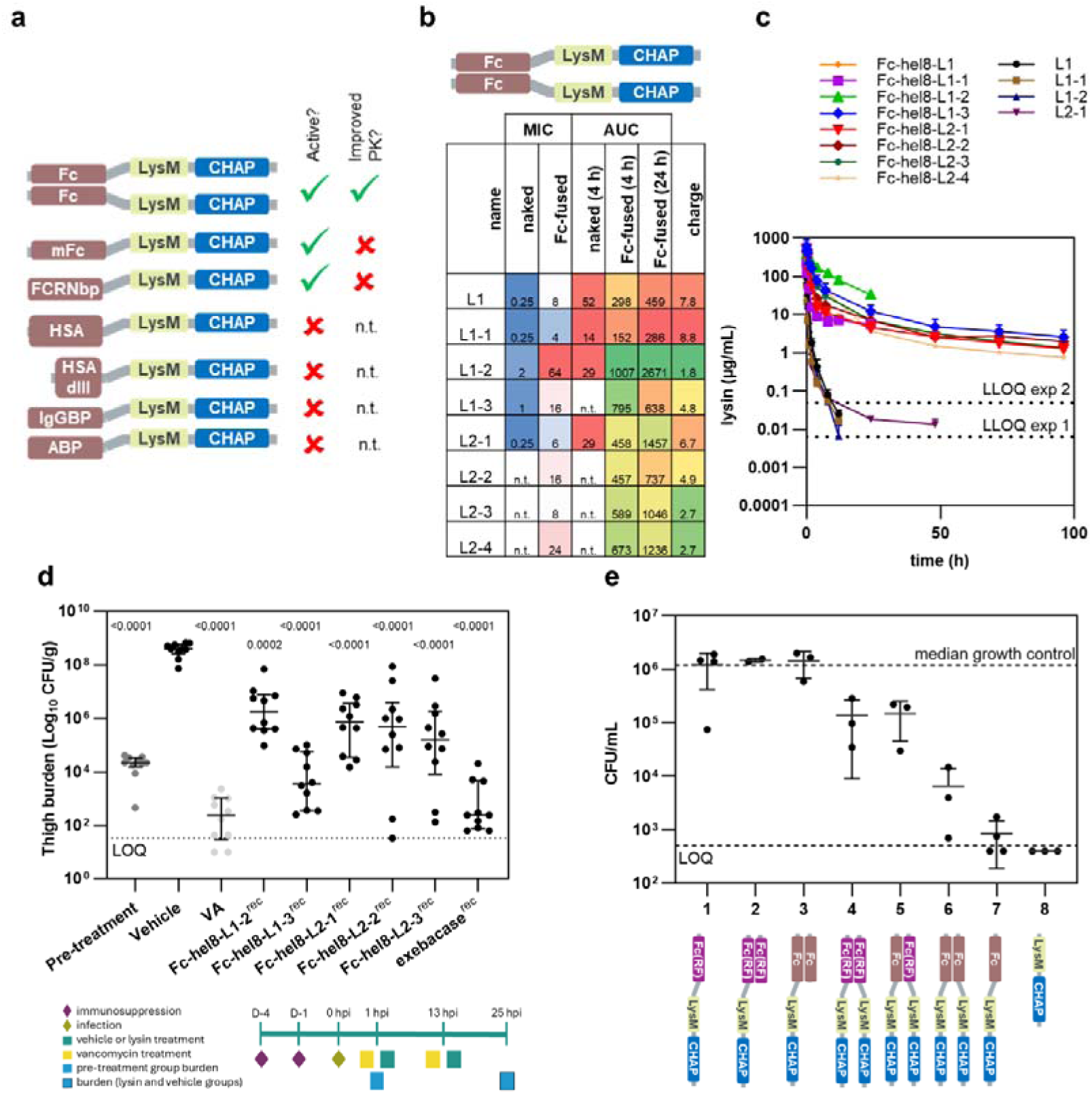
Selection of the Fc-LysM-CHAP architecture based on enhanced pharmacokinetics and in vivo efficacy. a.) Summary of pharmacokinetic (PK)-enhancing modalities fused to LysM-CHAP. Indicated PK-enhancing domains were fused to L1 and tested for growth inhibition against *Sa ATCC* 43300 in supernatant screens. Architectures with detectable activity were further purified and their PK-enhancing effect tested in mice. Fc: human IgG1-Fc domain; mFc: monomeric Fc domain; FcRNbp: peptide binder to neonatal Fc receptor; HSA: human serum albumin; HSAdIII: domain III of HSA; IgGbp: peptide binder to IgG1; ABP: peptide binder to HSA. b.) Activity and PK of naked and Fc-fused LysM-CHAP variants. Indicated LysM-CHAP variants were purified as naked lysins or fused to N-terminal Fc-hel8, followed by characterization of MIC (in µg/mL) against Sa ATCC 43300. AUC at 4 h and 24 h timepoints (AUC_4h_ and AUC_24h_; in µg*h*mL^-1^) were determined from PK curves shown in c.). Note that AUC_24h_ was only determined for Fc-fused lysins. n.t.: not tested; MIC: minimal inhibitory concentration; AUC: area under curve. c.) PK profiles of naked and Fc-fused LysM-CHAP variants. Purified lysins were injected as a single slow bolus intravenous injection via a lateral tail vein into C57BL/6J mice at a dose of 500 µg/mouse. Blood samples were drawn at indicated timepoints and lysin levels quantified by ELISA. Points on the graph represent a mean measured blood concentration of n=3 with standard deviation. Note that we show a composite of two separate experiments, with naked and Fc-fused derivatives of L1, L1-1, L1-2 (experiment 1; last measured timepoint at 24 h, lower limit of quantification (LLOQ) 0.00625 µg/mL) having been measured separately to naked and Fc-fused derivatives of L1-3, L2-1, L2-2, L2-3, and L2-4 (experiment 2; last measured timepoint at 96 h, LLOQ 0.05 µg/mL)). d.) Efficacy of lysins against Sa ATCC 43300 in a neutropenic mouse thigh infection model. Neutropenia was induced in male C57BL/6J mice (n=5) with cyclophosphamide and animals were infected with Sa ATCC 43300 by intramuscular injection of 5x10^3^ CFU/thigh in both thighs. Recombinant lysins, including exebacase, were applied at 1 hpi intravenously every 12 h (q12), at a dose of 500 µg/mouse. Vancomycin (VA) was used at dose of 100 mg/kg, q12 with treatment initiated 1 hpi. Data is depicted as log CFU/g of tissue in each thigh, harvested at 25 hpi (pre-treatment at 1 hpi). Each dot on the graph represents bacterial burden per gram of one thigh, with horizontal bars depicting median and interquartile range. Regular one-way ANOVA was used to calculate the significance compared to vehicle group with P values indicated in the graph. hpi: hours post infection; CFU: colony forming units; LOQ: limit of quantification (34 CFU/g). For graphical and statistical purposes, values under LOQ were assigned half of the LOQ value. e.) Bactericidal activity of different Fc-LysM-CHAP formats. Lytic activity of Fc-hel8-L1-3-variants with one or two LysM-CHAP arms, containing wild type Fc (brown) or Fc with RF mutations (to prevent Fc-binding to surface protein A; purple) and monomeric Fc variants thereof were tested in time kill assays in 80% human serum. Lysins were incubated at a final lysin concentration of 0.5 µM (refers to molar concentration of LysM-CHAP arms) with 10^6^ CFU/mL Sa ATCC 43300 for 3 h at 37 °C. Shown are individual replicates, mean and SD. Median growth control (untreated) is indicated as a dashed horizontal line. Activity of parental Fc-hel8-L1-3 is shown in column 6. LOQ: limit of quantification (500 CFU/mL). For graphical purposes, values under LOQ were assigned a value of 400 CFU/mL.

Next, we determined the PK and efficacy of Fc-LysM-CHAP recombinant protein (Fc-LysM-CHAP^rec^) variants in mice. Fc-LysM-CHAP^rec^ variants showed up to ∼90-fold increase in area under the curve (AUC) compared to their naked counterparts (Fig. 1b, c). The AUC inversely correlated with the overall charge (significant Pearson correlation; r=-0.8649, P=0.0056) (Fig. 1b and Extended Data Fig. 1c). In a mouse thigh infection model^24^, all Fc-LysM-CHAP^rec^versions were efficacious with notable differences in efficacy between the individual variants (Fig. 1d, 2.3- 4.9 log colony forming units (CFU)/mL reduction vs vehicle). Fc-hel8-L1-3 demonstrated greatest efficacy and was thus selected as the lead molecule for further optimizations.

To dissect the various features of the Fc-LysM-CHAP architecture, we investigated how size, valency and binding to Sa surface protein A (SpA) via the Fc^4^ affect in vitro potency. Systematic modifications of these properties were made by use of monomeric Fc variants^25^, hetero-dimeric Fc-hel8-L1-3 variants containing one lysin arm, or by introducing SpA-silencing (RF) mutations in the Fc^26^. Using time-kill assays we found that SpA binding and the bivalent engagement for binding and catalysis were beneficial for activity. Monomeric Fc-hel8-L1-3 showed the most promising in vitro potency but was excluded due to poor in vivo PK (Fig. 1e and Extended Data Fig. 1d). Taken together, these data pointed at cell surface or cell wall penetration and binding as key elements for further engineering and showed that dimeric Fc-hel8-L1-3 provides the most favourable balance between in vitro efficacy and PK.

### Sequence optimization

We hypothesized that Fc-hel8-L1-3 potency was limited by substrate access, binding and/or catalysis, and could be optimized through changes in linkers, modulation of cell binding properties and by mutagenesis of the catalytic pocket in the CHAP domain. We developed a screening platform to simultaneously record multiple parameters directly from supernatants in sufficiently high throughput to allow for rapid data-guided sequence optimization against these parameters (Extended Data Fig. 2a, Supplemental Discussion).

To structurally inform our optimization effort we performed size-exclusion chromatography small-angle X-ray scattering (SEC-SAXS) on Fc-hel8-L1-3. The analysis revealed an extended solution conformation of the protein with a maximum diameter and inter-CHAP distance of ∼30 nm (Fig. 2a, b). The finding of an extended conformation close to the dimensions of the Sa cell wall of ∼20-60 nm^27,28,29^ prompted us to systematically vary the length of the Fc-LysM linker (hel8 or linker-1) by changing the number of helical (EAAAK) repeats from 1 to 26. The longest (hel26) linker showed >4-fold improvement in Optical Density reduction rate (ODRR) and MIC relative to parental Fc-hel8-L1-3 (Fig. 2c and Extended Data Fig. 2b), but extension beyond hel14 reduced secretion levels to approximately half of those seen with Fc-hel8-L1-3, so hel14 was taken forward for optimization.

**Figure 2:**
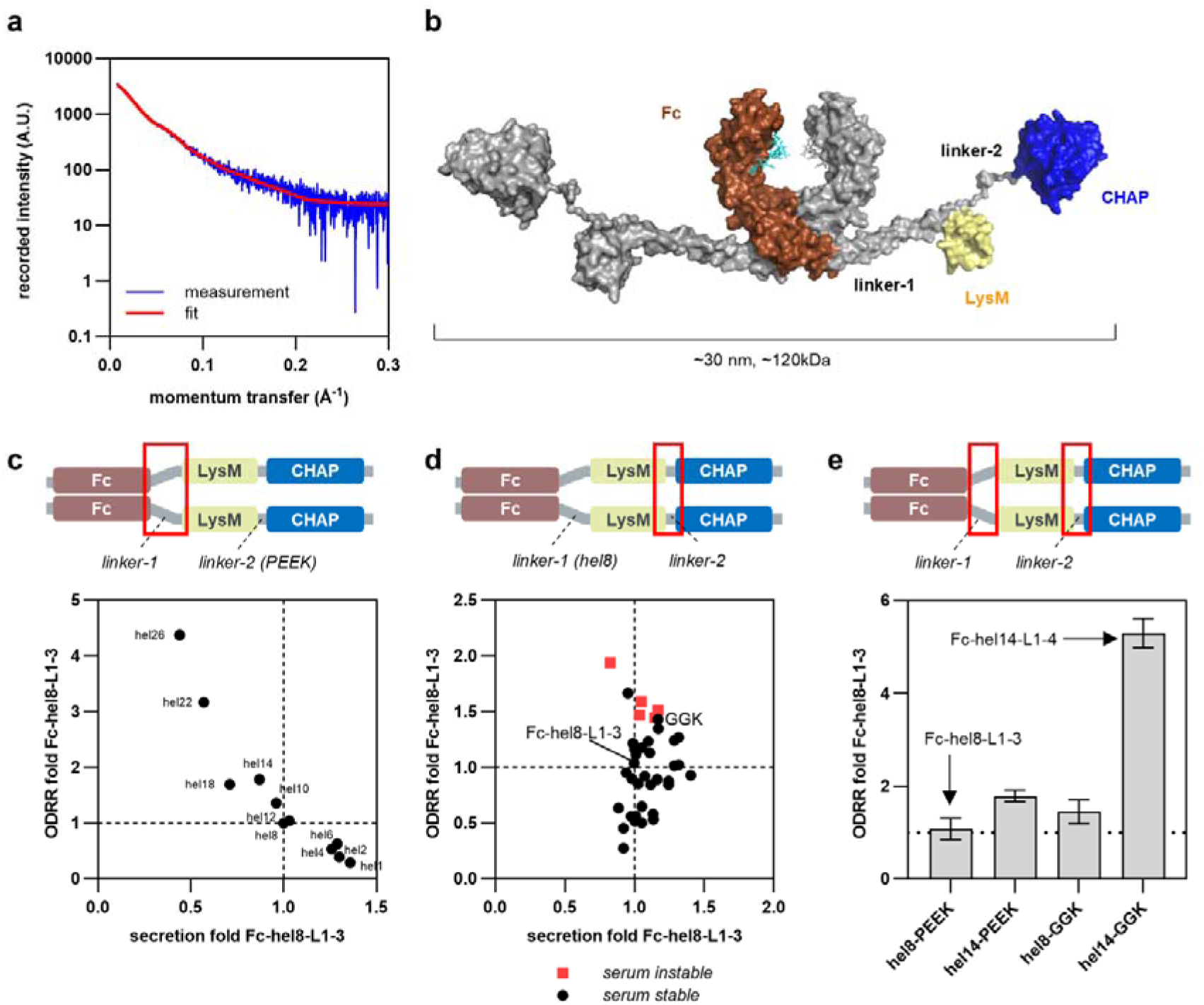
Linker optimization enhances lysin potency. a.) Size exclusion small angle X-ray scattering (SEC-SAXS) analysis of Fc-hel8-L1-3. SAXS scattering data overlaid with the theoretical scattering curves of the model shown under b.). A.U.: artificial units. b.) Solution model of Fc-hel8-L1-3 based on SEC-SAXS analysis. The model is shown in the surface representation. Fc (brown), LysM (yellow), CHAP (blue), linker-1 and linker-2 are indicated. Fc-glycans are shown as a stick model in turquoise. Note that the Fc-hinge was omitted from the modelling but contributes to dimer formation in Fc-hel8-L1-3. c.) Effect of helical linker length on lysin potency. Variants of Fc-hel8-L1-3 with truncations (hel6-hel1) or extensions (hel10-hel26) of linker-1 were transfected into Expi293 cells. Lysin secretion was determined by biolayer interferometry followed by measurement of activity using OD-reduction assays. OD reduction rates (ODRRs) and secretion levels were normalized against the parental Fc-hel8-L1-3 construct (containing a hel8 linker; indicated by vertical and horizontal dotted lines). d.) Effect of linker-2 amino acid sequence on lysin potency. As in c.) but with a set of Fc-hel8-L1-3 derivatives containing different sequences in the linker between the LysM and CHAP domains. Serum instable linkers are indicated in red. Secretion and activity of parental Fc-hel8-L1-3 is indicated by vertical and horizontal dotted lines, respectively. The shortlisted GGK linker is indicated. e.) Combination of hel14 and GGK in Fc-hel14-L1-4. ODRRs of Fc-hel8-L1-3 derivatives containing hel14, GGK or both hel14 and GGK were measured in supernatants. ODRRs were normalized to the parental Fc-hel8-L1-3 construct. Shown are means and standard deviations from at least two independent experiments.

For the LysM-CHAP linker (linker-2), 37 variants derived from LytN and its homologues were identified by InterPro searches and shortlisted for high-dimensional peptide feature similarity and predicted serum stability^30^. Linker-2 strongly impacted ODRR and MICs (0.25-2-fold and 0.5-6-fold, respectively, Fig. 2d and Extended Data Fig. 2c), suggesting that LysM and CHAP domains in FC-hel8-L1-3 require tight functional coordination for efficient bacterial killing. The “GGK” linker-2 was stable in human serum for >24 h (Extended Data Fig. 2d) and showed the highest improvement in ODRR and MIC. Combination of GGK with hel14 (Fc-hel14-L1-4) resulted in approximately 4-fold increased potency (Fig. 2e).

To gain insights into the mechanism of peptidoglycan hydrolysis and to guide further sequence optimization, we solved X-ray structures of the ligand-free apo-forms of the L1 and L2 CHAP domains as well as substrate-bound holo-forms of the L1 CHAP domain and of full length L1-3 (Fig. 3a, b, Extended Data Fig. 3a-f, Extended Data Table 3). The full-length structure of L1-3 shows that LysM interacts directly with the CHAP domain via a 275.4 Å^2^ interface and represents the first-ever visualization of a lysin CHAP domain in complex with an auxiliary targeting domain (Fig. 3a). In this conformation, the active site containing the catalytic Cys98 is occupied by the stem peptide (Fig. 3b) and aligns with a groove in LysM, which has been implicated in carbohydrate binding in other peptidoglycan hydrolases^31,32^. The continuous cleft formed by the two domains can accommodate a peptidoglycan fragment consisting of a MurNAc-GlcNAc chain and the stem peptide (Fig. 3a). Interestingly, the substrate is involved in interactions with both CHAP and LysM domains (Extended Data Fig. 3e). LysM-carbohydrate interactions and interactions between CHAP and the stem peptide are weak and transient (Extended Data Fig. 3g), as expected based on the observed interface bonds. The ternary complex of LysM, substrate and CHAP seen in our structure is thus likely formed by cooperative low-affinity interactions, explaining how LysM-CHAP can dissociate from its product after hydrolysis leading to processive engagement of the next peptidoglycan cross-link (see Supplemental Discussion and Extended Data Fig. 3h).

**Figure 3:**
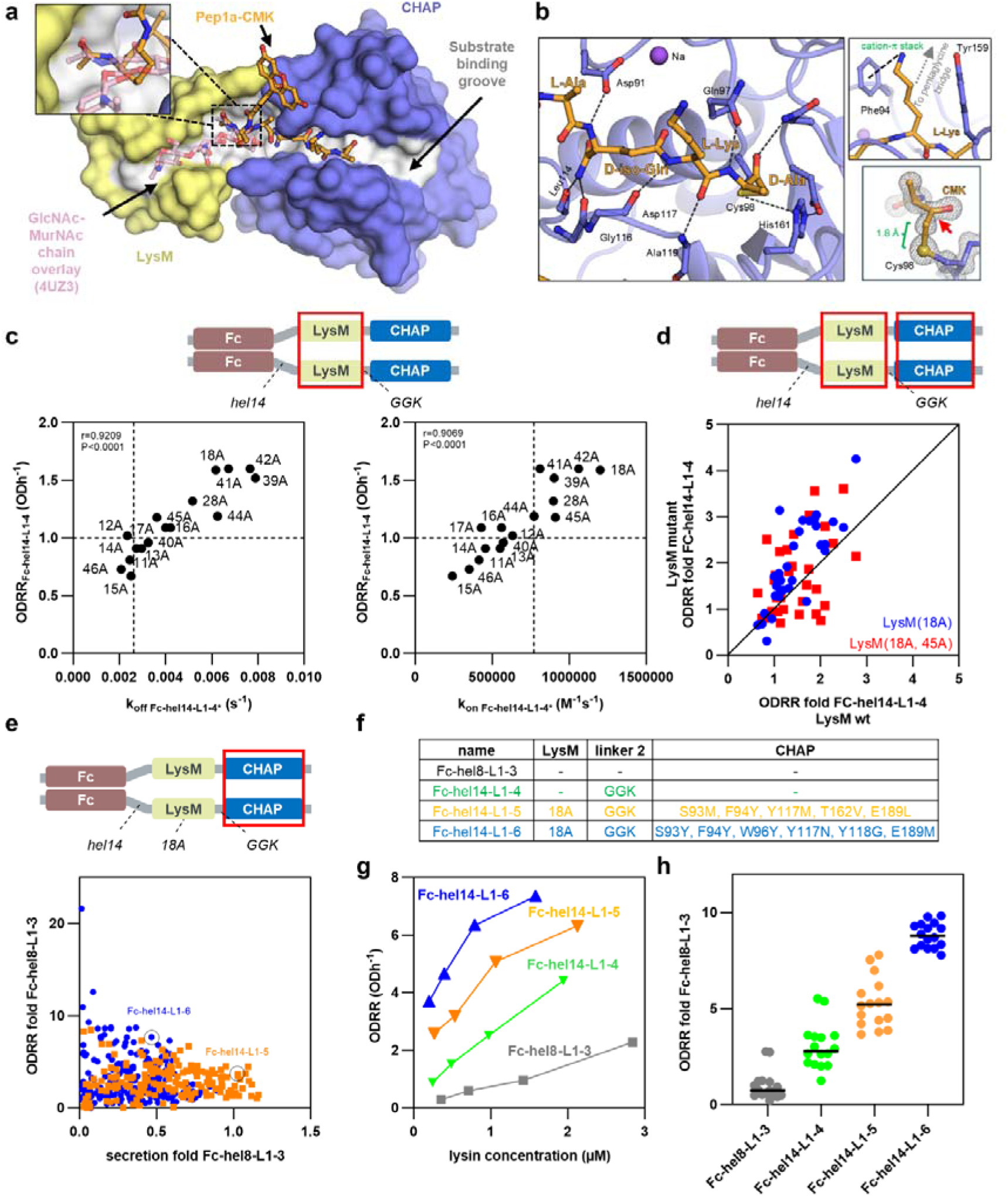
Structure-guided optimization of substrate recognition and catalysis by Fc-LysM-CHAP. a.) X-ray structure of LysM-CHAP L1-3 bound to a stem-peptide analogue (PDB: 11CI). Shown is a space-filling model of LysM (yellow) and CHAP (blue). The stem peptide analogue (orange) is shown as a ball-and-stick model. The modeled NAG-NAM chain from PDB:4UZ3 is shown as a transparent ball-and-stick model. The continuous substrate binding groove is indicated in grey. The inlet shows a zoomed-in view of the contacts between LysM and the stem-peptide analogue. b.) Interactions between stem peptide and the L1 CHAP active site (PDB: 11CH). The stem peptide is shown as a ball-and-stick model. Stem peptide amino acids are labeled in orange font. Amino acids from the CHAP domain in contact with the stem peptide are labeled in black font and shown as ball-and-stick models. Polar interactions between active site residues and the stem-peptide analogue are indicated with dotted lines. The top inlet shows the interactions between the stem peptide L-Lys residue and Phe94/Tyr159 in the CHAP domain (see Supplemental Discussion). The bottom inlet shows electron densities of the stem peptide and the catalytic Cys98 sulfur atom forming a 1.8 Å long covalent bond (indicated in green). The red arrow indicates the C_α_-atom of D-Ala in the stem peptide. c.) Correlation between Sa binding and lysin potency. Single alanine substitutions were introduced into the substrate binding pocket of the LysM domain in catalytically active Fc-hel14-L1-4 and catalytically inactive Fc-hel14-L1-4* (* denotes a Cys98Ser mutation in L1-4). Variants were purified from Expi293 supernatants. Catalytic rates of Sa (ATCC 43300) cell binding (k_on_) and dissociation (k_off_) of Fc-hel14-L1-4* derivatives were quantified using biolayer interferometry. ODRRs of catalytically active Fc-hel14-L1-4 counterparts were determined at 0.2 µM final enzyme concentration. k_off_ (left) and k_on_ (right) are plotted against ODRR of corresponding alanine mutants. Pearson coefficients (r) and P-values (P) are indicated for the correlation between k_off_ and ODRR and k_on_ and ODRR in the top left of each graph. d.) Differential impact of LysM mutations on the activity of CHAP variants. ODRRs of a set of 33 combinatorial motif engraftment (CME) designs of Fc-hel14-L1-4 containing either a wild type (wt) LysM domain or LysM domains harboring a Ser18Ala mutation (LysM(18A)) or Ser18Ala plus Ile45Ala mutations (LysM(18A,45A)) were determined in supernatants and normalized to the parental Fc-hel14-L1-4 construct. Rates of LysM(wt) variants are plotted against the corresponding LysM(18A) (blue) and LysM(18A,45A) (red) derivatives. e.) High throughput screen for active site mutants of Fc-hel14-L1-4(18A) with enhanced ODRRs. Active site variants of Fc-hel14-L1-4(18A) were generated using CME (blue) or inverse folding (orange) design strategies and transfected into Expi293 cells. Secretion levels in supernatants were determined by biolayer interferometry and lysin activities were determined by OD reduction assays. ODRRs were normalized to the parental Fc-hel14-L1-4 construct. Secretion levels were normalized to the original Fc-hel8-L1-3 construct. Secretion and ODRR were further expressed as fold changes relative to the parental Fc-hel8-L1-3 reference. Secretion and relative ODRRs of Fc-hel14-L1-5 and Fc-hel14-L1-6 are indicated. f.) Summary of engineering steps leading to Fc-hel14-L1-5 and Fc-hel14-L1-6. Substitutions in LysM, linker-2 and CHAP domain relative to Fc-hel8-L1-3 are indicated. g.) Dose-response behavior of lysin leads in OD reduction assays. Supernatants of Fc-hel8-L1-3, Fc-hel14-L1-4, Fc-hel14-L1-5 and Fc-hel14-L1-6 were serially diluted and measured for activity in OD reduction assays. Final assay lysin concentration (as determined by biolayer interferometry) is plotted against ODRR of each variant. h.) Summary of normalized ODRRs of lysin leads. ODRRs were recorded in supernatants from independent transfections and normalized relative to Fc-hel14-L1-4. ODRR were then expressed as fold changes relative to the Fc-hel8-L1-3 reference. Individual ODRRs are shown, horizontal lines indicate mean ODRRs.

Based on our X-ray data, we hypothesized that dimerization of the lysin in the Fc format leads to ‘too tight binding’, impairing product dissociation and hence catalytic activity. To reduce this binding affinity, we substituted positions 11-18, 28, 39, and 40-46 in the LysM carbohydrate binding channel to alanine, using the optimized Fc-hel14-L1-4 as background (Extended Data Fig. 3f). Catalytically active and inactive variants were expressed and purified and ODRRs and binding, respectively, were measured. For the latter, we determined k_off_ (dissociation rate constant) and k_on_ (association rate constant) to immobilized Sa cells by biolayer interferometry (BLI) (Extended Data Fig. 4a). ODRRs directly correlated with both k_off_ and k_on_ (Pearson correlation coefficients r=0.9209 and r=0.9069, respectively), indicating that rapid association and dissociation are needed for optimal potency (Fig. 3c). The LysM(18A) variant and a double mutant variant combining the most beneficial single mutations, LysM(18A,45A) scored highest in ODRRs (Extended Data Table 4) and were selected for follow-up engineering.

Next, we applied two approaches to engineer the enzymatic CHAP domain, based on the high-resolution structure of the L1 CHAP domain bound to the Sa stem peptide: 1. combinatorial motif engraftment (CME), which is based on co-occurring functional sequence motifs present in CHAP homologues, and 2. the use of InstaDeep’s DeepChain – a commercial AI platform – for large scale in silico sequence diversification of the CHAP catalytic pocket by inverse folding, followed by multidimensional ranking using optimized filters (Extended Data Fig. 2a).

In a pilot screen with 33 CME variants containing either LysM(wt), LysM(18A) or LysM(18A,45A), distinct CHAP variants were preferentially active in the context of specific LysM backgrounds, suggesting tight coupling between peptidoglycan binding and hydrolysis (Fig. 3d, Extended Data Fig. 4b-d). CME variants harbouring LysM(18A) achieved the highest improvements in ODRR and MIC, so LysM(18A) was chosen as the backbone for further engineering in the Fc-hel14-L1-4 background.

We next screened a larger library of 225 active site variants generated by CME and of 175 variants generated by inverse folding. The combined sequences displayed improvements in ODRRs up to 22-fold (Fig. 3e) and MICs up to 20-fold vs Fc-hel8-L1-3 (Extended Data Fig. 4e). We observed a negative correlation between lysin activity and secretion among the most active CHAP derivatives, potentially explained by the destabilizing effect of function-enhancing mutations reported during the evolution of many enzymes^34,35^. Interestingly, this trade-off between secretion and activity was most pronounced in the CME library, in which functional motifs had been grafted into the catalytic pocket without introducing compensating mutations. In contrast, mutations designed by inverse folding were better tolerated, albeit at the cost of smaller increases in potency (Extended Data Fig. 4f).

Variants were shortlisted if they showed meaningful improvements in at least two of the three activity dimensions that had been measured (i.e. ODRR > 5x and/or MIC > 8x improved over Fc-hel8-L1-3 against either tested Sa strain). Shortlisted variants were purified and compared for biophysical and biochemical properties and in growth inhibition and time kill assays. CME variant Fc-hel14-L1-6 and inverse folding variant Fc-hel14-L1-5 exhibited optimal properties after purification and were selected as lead candidates (Fig. 3f). Across multiple supernatant screens, they displayed 10-20-fold improved dose-response behaviour in OD reduction assays, 5-9 times increased ODRR (Fig. 3 g, h) and up to 16-fold reduced MICs versus parental Fc-hel8-L1-3 (Extended Data Fig. 4g).

### In vitro potency and synergy with antibiotics

We investigated the dose-response behaviour of Fc-hel8-L1-3 and engineered variants alone or in combination with vancomycin, an antibiotic for the treatment of severe and life-threatening methicillin-resistant Sa (MRSA) infections^36^ (Fig. 4a). For all tested LysM-CHAP lysins, synergy with vancomycin was observed across the lysin concentration series, indicating a 4-8-fold increase in potency upon combination of each lysin with vancomycin (Fractional eradication concentration index, FECI = 0.06). Synergy between Fc-hel14-L1-5 or Fc-hel14-L1-6 with vancomycin was observed across multiple staphylococcal species (FECI=0.06) (Extended Data Fig. 5a) and at different vancomycin concentrations (Extended Data Fig. 5b; Supplemental Discussions). Co-treatment with lysin and vancomycin reduced the median minimal eradication concentration (MEC) by 4-8-fold compared with lysin alone; Fc-hel14-L1-6 and Fc-hel14-L1-5 showed 8-16-fold lower MEC vs Fc-hel8-L1-3, both in presence and absence of vancomycin (Fig. 4b). The addition of vancomycin had a less pronounced impact on MEC (1-4 fold) for exebacase, lysostaphin and non-Fc tagged LysM-CHAPs (Extended Data Fig. 5c). In the presence of vancomycin (16 µg/mL), the MEC in human serum against Sa BAA1717 for Fc-hel14-L1-6 (87 nM) was similar to that of lysostaphin (57 nM) (Extended Data Fig. 5c).

**Figure 4:**
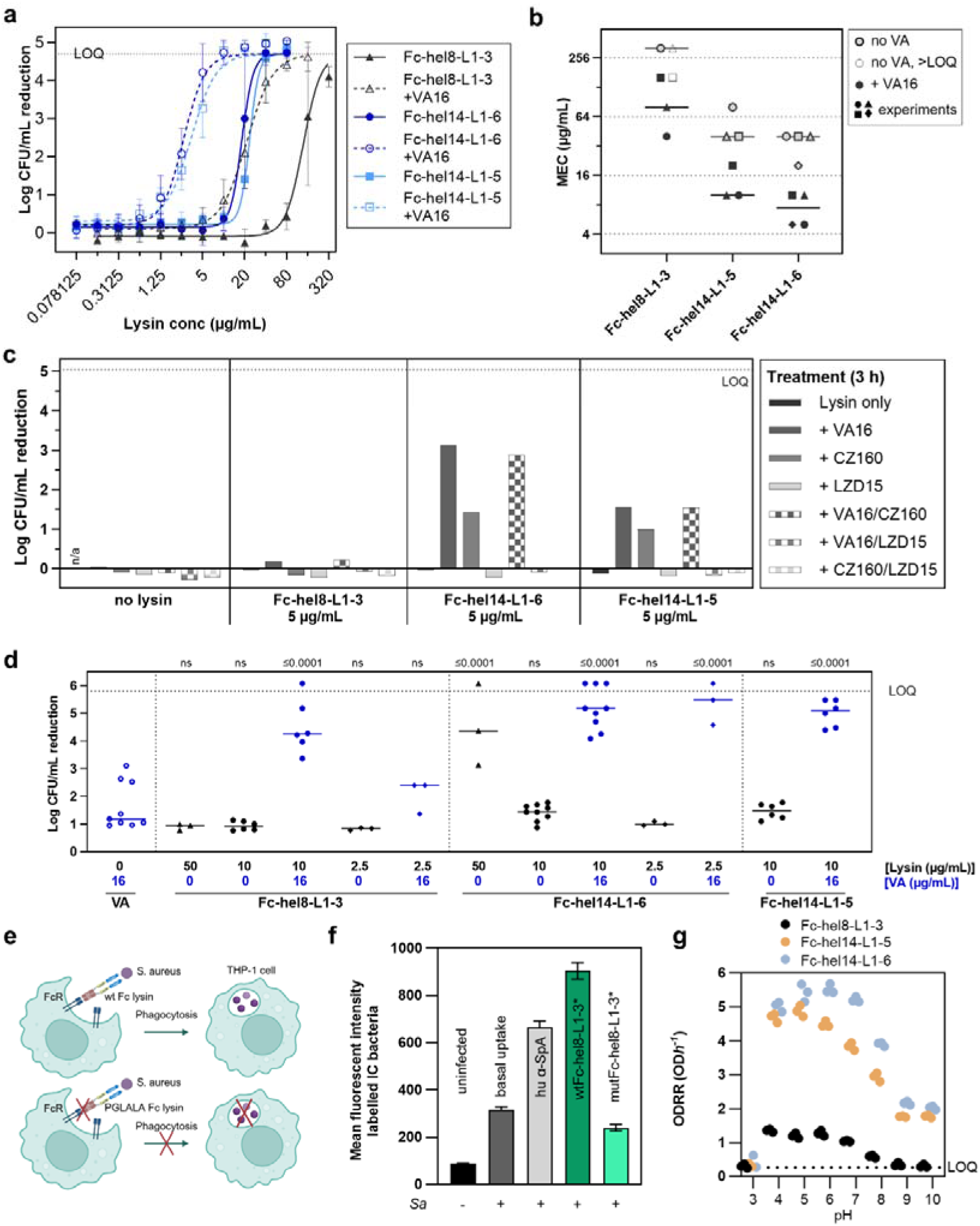
Synergy with cell wall–active antibiotics enhances efficacy of Fc–LysM17CHAP lysins. a.) 4-8 fold increased potency upon lysin combination with vancomycin. Lysin dose-response series were tested for 3 h in 80% human serum with or without 16 µg/mL vancomycin on Sa BAA-1717. Mean with SD of three independent experiments is shown. Median LOQ (eradication to below limit of quantification (blq) is indicated as dashed horizontal line. The Four-Parameter Logistic (4PL) curves were fitted to the data with the top constrained to median LOQ. b.) Minimal eradication concentration (MEC) of lysins with or without vancomycin on Sa BAA-1717 in 80% human serum. Concentration series of lysins were tested on 5×10^7^ to 2×10^8^ CFU/mL Sa cells as single (grey symbols) or combination treatment with vancomycin at fixed concentration of 16 µg/mL (black symbols). MECs above the highest tested concentration are shown as empty symbols. Each symbol represents an independent MEC determination, median is indicated. c.) Activity of lysins with or without antibiotic co-treatment. Fc-tagged lysins (5 µg/mL) were tested alone or in combination with different antibiotics in a time kill assay on Sa BAA-1717 for 3 h in 80% human serum in technical duplicates. Concentrations (µg/mL) of antibiotics (vancomycin: VA, cefazolin: CZ and linezolid: LZN) are stated in the legend. Log_10_ CFU/mL reduction was calculated relative to the buffer-treated control. d.) Reduction of Sa BAA-1717 from in vitro simulated endocardial vegetations. Simulated endocardial vegetations (clots) were incubated for 24 h with stated lysins at 50 µg/mL (triangles), 10 µg/mL (closed circles), 2.5 µg/mL (diamonds) and antibiotic alone (open circles) with (blue) or without (black) 16 µg/mL vancomycin. Log CFU/mL reduction was calculated relative to the untreated control. Replicates from different experiments with median are shown. Ordinary one-way ANOVA (P<0.0001) was performed on log-transformed values followed by Dunnett’s multiple comparison test against vancomycin-treated clots (open circles). P values are indicated in the graph; ns, not significant. e.) Schematic representation of Fc-lysin mediated uptake of bacteria by human THP-1 cells. Lysins containing PGLALA mutation in their Fc domain are unable to interact with Fc receptors on immune cells and thus do not lead to an increase of bacterial uptake compared to the basal uptake. (Scheme drawn using BioRender.) f.) Uptake of Sa BAA-1717 by human THP-1 monocytes in presence of lysins and control human anti-SpA antibody (hu α-SpA; 100 nM). THP-1 cells were exposed to fluorescently labeled (carboxyfluorescein succinimidyl ester – CFSE) bacteria at a multiplicity of infection (MOI) of 10 (basal uptake) in presence of enzymatically inactive lysins containing either wt Fc (wtFc-hel8-L1-3) or inert, PGLALA modified Fc (mutFc-hel8-L1-3) for 1 h. The cells were then washed, fixed and fluorescence was measured using flow cytometry. The bars represent technical triplicate with standard deviation. g.) Activity of Fc-LysM-CHAP variants at different pHs. ODRRs of Fc-hel8-L1-3, Fc-hel14-L1-5 and Fc-hel14-L1-6 were measured at a final enzyme concentration of 0.5 μM and in different buffers with pHlllranging from 3 to 10. LOQ: limit of quantification at 0.27 ODh^-1^.

We also tested other classes of antibiotics which are typically used in the clinic to treat bacteraemia and endocarditis, including β-lactams, lipopeptides, and oxazolidinones. Little activity was observed in a time kill assay against Sa BAA-1717 with lysins alone at 5 µg/mL or antibiotics alone at concentrations corresponding to 8-32-fold MIC (Fig. 4c). In contrast, combination treatments with both vancomycin and cefazolin (both cell wall active antibiotics^37^) increased the log_10_ CFU/mL reduction from essentially 0 for the lysin alone to 2-3, indicating strong synergy (Fig. 4c). Combining vancomycin and cefazolin with any of the Fc-LysM-CHAP variants did not increase the reduction in bacterial load compared to vancomycin only plus with lysin. Interestingly, addition of the protein synthesis inhibitor linezolid to either vancomycin or cefazolin and Fc-LysM-CHAP showed antagonistic effects compared to the co-treatments described above (Fig. 4c).

To mimic in vivo settings, such as those encountered in staphylococcal infective endocarditis, an ex vivo model of simulated endocardial vegetations was developed, using fibrin clots generated in the presence of human plasma and thrombin^38^, in which bacteria persist in a biofilm-like state^2^. In this model, vancomycin (16 µg/mL) or the Fc-LysM-CHAP variants (10 µg/mL) showed negligible or low activity (<1.5 log CFU/mL reduction), but their combination was highly effective (4–5.5 log CFU/mL reduction) (Fig. 4d). Fc-hel14-L1-6 was active at the lowest tested concentration of 2.5 µg/mL in presence of vancomycin but a dose of 50 ug/mL was required to achieve the same activity as monotherapy, indicating that vancomycin decreases the MEC of Fc-hel14-L1-6 by ≥20-fold.

The fusion of lysins to Fc does not only extend the in vivo half-life but can, in principle, also impart additional mechanisms of action to the fusion protein. Fc-mediated phagocytosis using the lysin binding module as a bacterial surface anchor has been demonstrated for lysibodies^39^. The survival of Sa inside phagocytic cells was shown to be critical to bacterial persistence and tissue distribution in a mouse sepsis model with delayed treatment onset. In this delayed treatment model, an antibody-antibiotic conjugate, which releases the antibiotic intracellularly, demonstrated substantially better efficacy than the extracellular antibiotic^40^. In our study, antibody-dependent cellular phagocytosis into human THP-1 monocytes was observed for Fc-hel8-L1-3 at similar levels observed with a specific anti-Sa antibody (directed against SpA) but not for its Fc-silenced counterpart (PGLALA^42^) (Fig. 4e-f). This Fc-variant did not change the in vitro potency or the biophysical properties of Fc-hel8-L1-3. Moreover, the Fc-LysM-CHAP variants were fully active at pH conditions mimicking the acidified phagolysosome (pH 4-7; Fig. 4g).

### PK and efficacy studies

To test the in vivo efficacy of the lead lysin candidates, an extended (72 hour [h] instead of 24 h) murine Sa sepsis model was developed to better mimic the clinical situation and to capture the treatment effect on intracellular bacteria (Sa proliferates in the intracellular niche only >7 hours post-infection [hpi]^40^). In a first cohort of mice, treatment was started at 1 hpi. Fc-hel14-L1-6^rec^ (3.5 mg/kg, q12h) led to significant log CFU reduction in the kidneys vs vehicle (P=0.0130), as did vancomycin at 25 mg/kg (P=0.0066) (Fig. 5a). In a second cohort of mice, treatment was started after an initial bacteraemia establishing phase at 13 hpi. Vancomycin showed significant efficacy vs vehicle (P<0.0001) but could not reduce the bacterial burden below stasis (P>0.9999). The combinations of Fc-hel14-L1-6^rec^ or Fc-hel14-L1-5^rec^ with vancomycin, but not single treatments (or Fc-hel8-L1-3^rec^/ exebacase plus vancomycin), led to a significant reduction of the median bacterial burden vs stasis by ∼2 log10 units (P=0.0023 and 0.0167 respectively; Fig. 5b).

**Figure 5:**
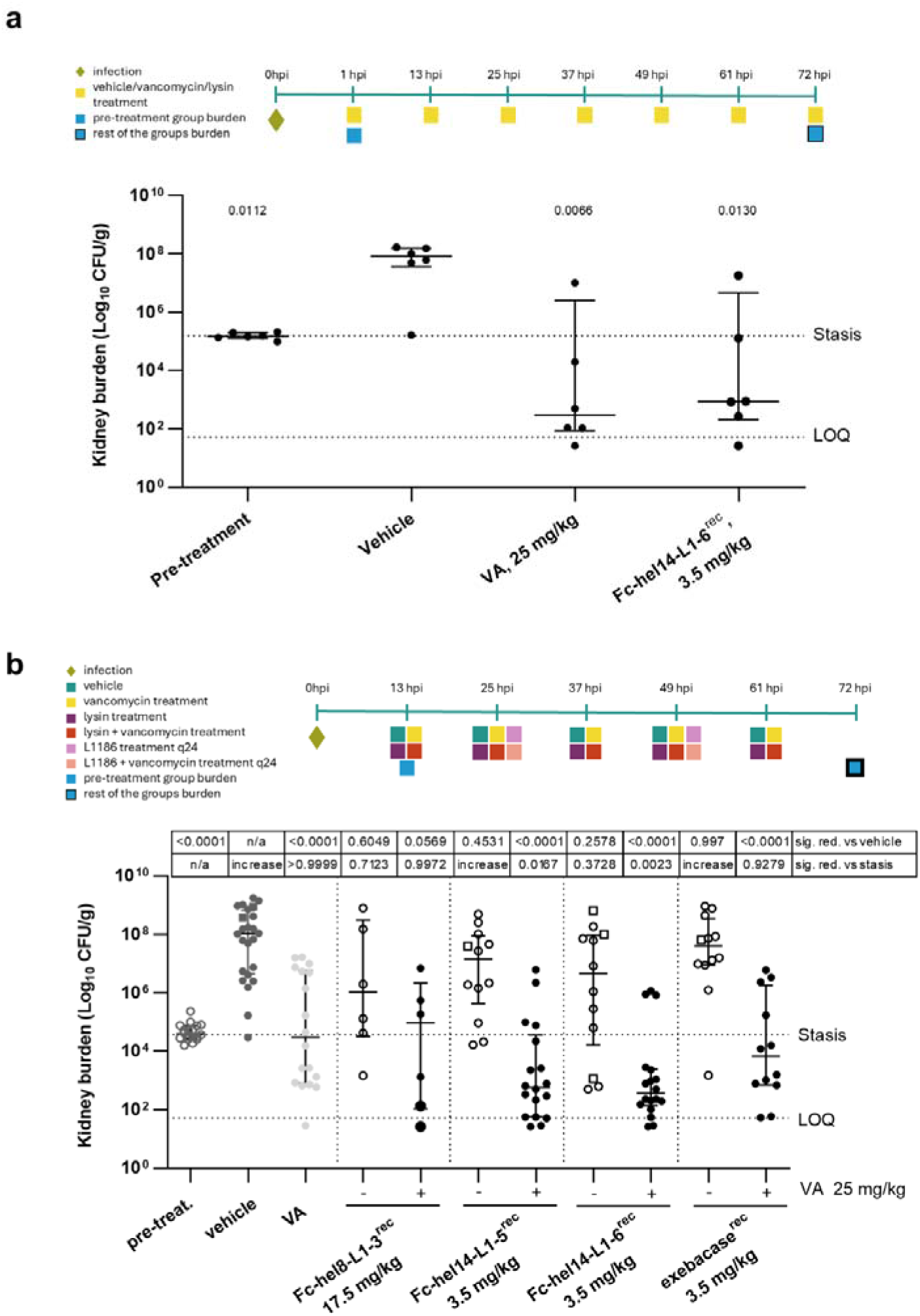
Combination therapy with Fc–LysM11CHAP lysins improves outcomes in a delayed11treatment sepsis model. a.) Efficacy of Fc-hel14-L1-6 and Vancomycin in a CD1 mouse Sa BAA-1717 intravenous (IV) sepsis model. Both therapies were administered at 1 hpi, IV, every 12 h (q12), Fc-hel14-L1-6 at the dose of 3.5 mg/kg and vancomycin (VA) at 25 mg/kg. Data is depicted as log CFU/g of tissue in kidneys, harvested at 72 hpi (1 hpi for pre-treatment group). Points on the graph represent CFU counts in individual mice (n=6), with horizontal lines indicating median CFU burden and interquartile range. Brown-Forsythe ANOVA with Welch’s correction was used to calculate the significance compared to vehicle group on log transformed data, with P values indicated in the graph. hpi: hours post infection; CFU: colony forming units; LOQ: limit of quantification at 52 CFU/g; For graphical and statistical purposes, values under LOQ were assigned half of the LOQ value. b.) Efficacy of Fc-hel8-L1-3, Fc-hel14-L1-6, Fc-hel14-L1-5, exebacase, and Vancomycin, as mono- or combination treatments in a CD1 mouse Sa BAA-1717 intravenous (IV) sepsis model. Treatments were initiated at 13 hpi. Fc-hel14-L1-6^rec^, Fc-hel14-L1-5 ^rec^, and exebacase ^rec^ were dosed at 3.5 mg/kg, IV, every 12 h (q12h), while Fc-hel8-L1-3 ^rec^ was dosed at 17.5 mg/kg, IV, q24. Vancomycin (VA) was applied at 25 mg/kg, subcutaneously, q12. Data is depicted as log CFU/g of tissue in kidneys, harvested at 72 hpi (13 hpi for pre-treatment group). Points on the graph represent CFU counts in individual mice, with horizontal lines indicating median CFU burden and interquartile range. Squares indicate mice that were terminated before the end of the study due to reaching clinical endpoints. Brown-Forsythe ANOVA with Welch’s correction was used on log transformed data to calculate the significance compared to vehicle or stasis (pre-treatment) groups with P values indicated in the table above the graph, for the groups where reduction in bacterial burden was detected. Significant increase was labeled as “increase”. hpi: hours post infection; CFU: colony forming units; LOQ: limit of quantification at 52 CFU/g; For graphical and statistical purposes, values under LOQ were assigned half of the LOQ value.

We next investigated if LNP-encapsulated mRNA encoding for Fc-hel8-L1-3 (Fc-hel8-L1-3^RNA^) was able to produce active lysin in situ in mice. The mRNA-LNP used had been previously optimized for secretion and translation in the liver, as well as for reduced immunogenicity by introducing modified nucleosides^43,44^. The PK of Fc-hel8-L1-3^RNA^ was determined for two LNP-mRNA dose levels (2.0 and 0.5 mg/kg) and compared to that of Fc-hel8-L1-3^rec^ (25 mg/kg) upon intravenous (IV) injection in C57Bl/6 mice. The AUC for the 0.5 mg/kg Fc-hel8-L1-3^RNA^ cohort was in the range of Fc-hel8-L1-3^rec^ and the Cmax values (440 and 60 ug/mL, at the high and low LNP-mRNA doses, respectively), as well as the Tmax (∼8 h) (Fig. 6a) were in the ranges observed for LNP-mRNA-delivered monoclonal antibodies^45,46^. The time over MIC increased to 48 – 96 h for Fc-hel8-L1-3^RNA^ vs <24 h for Fc-hel8-L1-3^rec^ (as expected based on the continuous translation of protein over 24-48 h^47^). We then determined the efficacy of Fc-hel8-L1-3^RNA^ in the Sa murine sepsis model at a single dose of 2.0 mg/kg and compared it to the Fc(PGLALA) variant with silenced Fc effector function^48^ (Fc(PGLALA)-hel8-L1-3^RNA^). Both treatments were highly efficacious and on par with multiple doses of vancomycin (Fig. 6b), indicating that active lysin is produced in situ and that its efficacy is not solely driven by the Fc function. In a second experiment, we interrogated the efficacy of Fc-hel14-L1-6^RNA^ at dose levels of 0.2, 0.5 and 2.0 mg/kg mRNA-LNP and of Fc(PGLALA)-hel14-L1-6^RNA^ at the intermediate dose level of 0.5 mg/kg. Fc-hel14-L1-6^RNA^ led to significant efficacy in the kidneys at all dose levels (∼3, 4 and 5 log CFU reduction vs vehicle with P=0.0298, P=0.0108 and P<0.0001 respectively, Fig. 6c) whereas Fc(PGLALA)-hel14-L1-6^RNA^ was ineffective (P=0.5552), indicating that the Fc function is partly responsible for in vivo efficacy.

**Figure 6:**
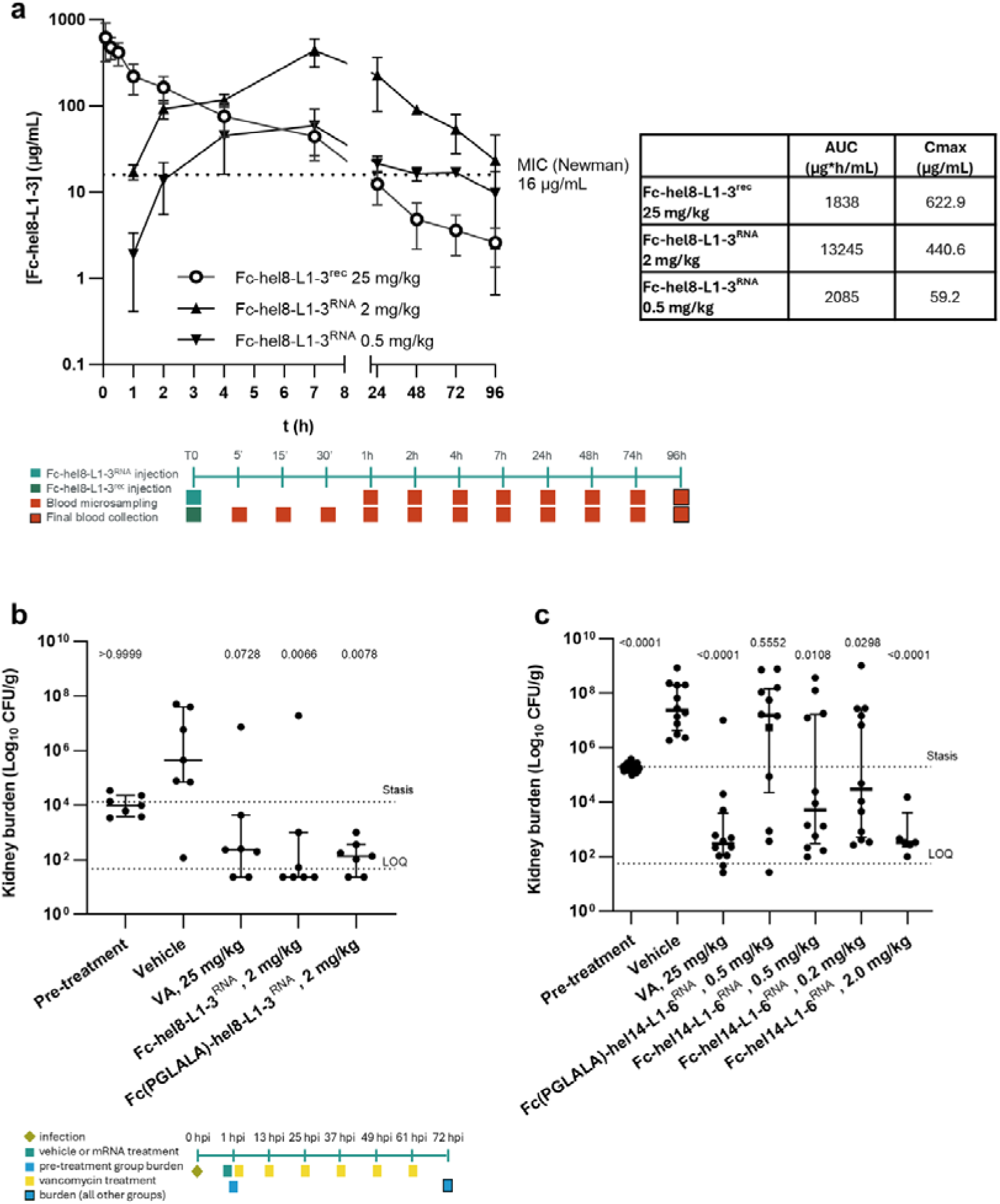
RiboLysin delivery enables in situ production and in vivo efficacy of Fc11LysM11CHAP lysins. a.) Pharmacokinetic (PK) profiles of Fc-hel8-L1-3 (2.0 and 0.5 mg/kg) and Fc-hel8-L1-3 (25 mg/kg). Test articles were administered at the indicated dose as a single slow bolus intravenous injection via a lateral tail vein on C57Bl/6J mice. Points on the graph represent a mean measured concentration of n=3 with standard deviation. b.) Efficacy of Fc-hel8-L1-3^RNA^, Fc-hel14-L1-6^RNA^ and their Fc-inert (PGLALA) variants in a CD1 mouse Sa NCTC 8178 intravenous (IV) sepsis model. Test articles were administered at a dose of 2 mg/kg, at 1 hpi as a single slow bolus IV injection via a lateral tail vein. Vancomycin (VA) was used as a comparator drug at dose of 25 mg/kg, IV, every 12 h (q12). Data is depicted as log CFU/g of tissue in kidneys, harvested at 72 hpi (1 hpi for pre-treatment group). Points on the graph represent CFU counts in individual mice (n=7), with horizontal lines indicating median CFU burden and interquartile range. Kruskal-Wallis test was used to calculate the significance compared to the vehicle group on log transformed data, with numbers indicating the P values. hpi: hours post infection; CFU: colony forming units; LOQ: limit of quantification at 47 CFU/g; For graphical and statistical purposes, values under LOQ were assigned half of the LOQ value. c.) Efficacy of Fc-hel14-L1-6^RNA^ and its Fc-inert (PGLALA) variant in a CD1 mouse Sa BAA-1717 intravenous (IV) sepsis model. All test articles were administered at 1 hpi as a single slow bolus IV injection via a lateral tail vein at dose of 0.2 mg/kg, 0.5 mg/kg, or 2 mg/kg, as indicated in the graph. Vancomycin was used as a comparator drug at dose of 25 mg/kg, IV, every 12 h (q12). Data is depicted as log CFU/g of tissue in kidneys, harvested at 72 hpi (1 hpi for pre-treatment group). Points on the graph represent CFU counts in individual mice (n=6-24), with horizontal lines indicating median CFU burden and interquartile range. Squares indicate mice that were terminated before the end of the study due to reaching clinical endpoints. Brown-Forsythe ANOVA with Welch’s correction was used to calculate the significance compared to the vehicle group on log transformed data, with numbers indicating the P values. hpi: hours post infection; CFU: colony forming units; LOQ: limit of quantification at 56 CFU/g. For graphical and statistical purposes, values under LOQ were assigned half of the LOQ value.

## Discussion

In summary, we show that following multi-dimensional protein optimization efforts, we converted an endogenous cell wall remodelling enzyme into the potent anti-staphylococcal therapeutic candidate Fc-hel14-L1-6. This candidate targets both intra- and extracellular bacteria as well as bacterial vegetations and strongly synergizes with cell wall active antibiotics. Fc-hel14-L1-6 demonstrates a clear advantage over the standard of care (SOC) and previous clinical lysin candidates in preclinical models. When delivered from mRNA, the engineered lysin has in situ expression and half-lives comparable to monoclonal antibodies^46^.

Dimerization of the lysin in the Fc architecture caused a reduction in potency, which could be rescued by modulation of binding kinetics, active site engineering and optimization of the Fc-lysin linker length and sequence. We hypothesize that a sufficiently long linker is required to provide access to deeper peptidoglycan layers (∼ 40 nm CHAP to CHAP for hel14 is in the same range as the peptidoglycan thickness) and additional degrees of freedom for the processing of the three-dimensional substrate. We additionally found that lysin charge has dual effect: increased charge is beneficial for activity, but detrimental for PK.

Using these design principles, we show that an Fc-functionalized lysin (here LysM-CHAP) is a feasible, versatile modality to target Sa in clinically relevant infection models, combining lysin and Fc modes of action, and a promising RiboLysin candidate. Fc-engineering can be further exploited to improve half-life^49^ and potentially reduce immunogenicity^50^.

The RiboLysin design space and development path described here can be adapted for lysins targeting other pathogens, to streamline the R&D process for novel countermeasures against emerging bacterial threats.

## Data availability

Structural data from X-ray crystallography has been deposited in the RCSB Protein Data Bank (PDB) under the accession numbers 11CF, 11CG, 11CH and 11CI.

SAXS raw data has been submitted for deposition in the SASBDB (accession code: SASDZ46).

## Code availability

Code for the foundational inverse folding models used in this study, LigandMPNN and protein DPO, is publicly available and detailed in their respective original publications cited in the Materials and Methods. DeepChain is an industry-grade commercial AI platform developed by InstaDeep and is available at https://deepchain.bio. A detailed algorithmic workflow describing the in silico sequence diversification, multidimensional ranking, and optimized filtering parameters is provided in the Materials and Methods. Custom scripts or specific parameters used for the geometric constraint filtering and the activity predictor are available from the corresponding author upon reasonable request. Demonstrations and trial access to the DeepChain platform are available from the developer or the corresponding author upon reasonable request.

## Supporting information

Supplemental Discussion

## Acknowledgements

We thank Drs. Michele Mutti and Karsten Krey for insightful discussions and scientific input on the screening strategies. We are especially grateful to Gurli Wertanek, Manuel Maluenda and Sruthi Kurugodu for their excellent technical work. We thank Anna-Margarita Schötta for performing yeast-display experiments. We thank Drs. Alexander Muik, Alibek Galeev, Anna Spier, Fiona Powell and Amanda Gallagher for critical review of the manuscript, Drs. Ilya Kizhvatov and Maren Lang for the development of protein engineering scripts using the Rosetta algorithms, Drs. Johannes Kraml and Olaf Otherson for providing essential insights into the mechanism of peptidoglycan hydrolysis using molecular dynamics simulations and Drs. Karsten Schnatbaum and Michael Drosch at JPT peptide technologies for the synthesis of pep1a-CMK. Also, we would like to acknowledge Acuitas Therapeutics for LNP technology and thank the teams at Acuitas as well as the formulation department at BioNTech SE for providing us with the mRNA-LNP material used in this study. We acknowledge Evotec UK (Ltd) for performing the animal studies, the Vienna BioCenter core facilities (Protein Technologies) for protein production and BioSAXS for the recording and analysis of SEC-SAXS data of Fc-hel8-L1-3.

## Author information Authors and Affiliations

Benedikt Bauer and Zehra Visram contributed equally to the work

BioNTech R&D Austria

Adriana Badarau, Benedikt Bauer, Zehra Visram, Renping Qiao, Andrea Majoros-Hashempour,, Svetlana Durica-Mitic, Robert Kluj, Donaat Kestemont, Ann-Katrin Kieninger, Monika Kulig, Rocio Berdaguer, Timo Schwebs, Manuel Zerbs, Philipp Czermak, Marina von Freyberg, Julia Schmidt, Laura Schmidberger, Dominik Mayer, Lorenzo Corsini

Former employees of BioNTech R&D Austria Johannes Söllner, Orla Dunne, Maria Protano, BioNTech US

Hannah Miller, Gavin Palowitch, Charles Dulberger, Maria Massaro InstaDeep

Nikita Ivanisenko, Hippolyte Jacomet, Didier Ngatcha Bakoue, Gali Sela

Former employees of InstaDeep Francesco Oteri

Corresponding authors

Adriana Badarau, Lorenzo Corsini

## Contributions

LC conceptualized and directed the RiboLysin project overall, reviewed data and progress, AB conceptualized the Fc-lysin modality, directed the research and led the project. BB and ZV contributed equally to the work, BB directed protein engineering and ZV directed the in vitro functional characterization and in vivo work. AB, BB, ZV, RQ, JS, LC, designed research and provided supervision. BB, ZV, RQ, AMH, SDM, OD, RK, DK, AK, MK, RB, TS, MZ, PC, MF, JSC, LS, DM, MP performed the experiments. AB, BB, ZV, RQ, AMH, SDM, OD, RK, DK, AK, MK, RB, TS, MZ, PC, MF, JSC, LS, DM, MP analyzed and interpreted the data. HM, GP, CD, MM performed the crystallography and X-ray structure determination work. JS, NI, HJ, DNB, FO and GS performed the AI-guided CHAP optimizations. AB, BB and ZV wrote the manuscript with input from RQ, AK and AMH. All authors reviewed the manuscript.

## Ethics declarations Competing interests

AB, BB, ZV, RQ, AMH, SDM, RK, DK, AK, MK, RB, TS, MZ, PC, MF, JSC LS, DM, HM, GP, CD, MM, LC are employees of BioNTech and hold shares in the company. JS, OD and MP are former employees of BioNTech and hold shares in the company. NI, HJ, DNB, GS are employees of InstaDeep, a subsidiary of BioNTech. HJ and DNB hold shares in BioNTech. FO is a former employee of InstaDeep and holds shares in BioNTech. AB, BB, ZV, RQ, SDM, RK, DK, JS, OD, LC, NI, HJ, DNB, GS and FO are inventors on patent applications related to the technology described in the manuscript.

## Methods and Materials

### Directed evolution of LysM-CHAP

Directed evolution of L0 was performed using Pichia pastoris (Pp) strain GS115 in the following way: The L0 sequence and 18 homologues of L0 were codon optimized for expression in Pp in a way that all identical amino acids between the variants were encoded by the same DNA sequence. DNA fragments from each of the homologues and L0 were recombined by DNA shuffling^1^ and then cloned into a coupled Pp display/secretion vector derived from pPIC9K by Gibson Assembly^2^. The Pp display/secretion vector contained the following elements in the listed order: a methanol-inducible AOX1 promoter, a pre-OST signal sequence from S.cerevisiae (Sc), the pro sequence of alpha mating factor from Sc, an HA-epitope tag, the LysM-CHAP sequence, an F2A ribosome skipping peptide^3^, a V5 epitope tag, and the C-terminal 320 amino acid fragment of Sc Sag1p. The pPIC9K vector was further modified to contain an autonomous replicating sequence^4^. The vector was designed in a way that upon methanol induction, roughly 50% of the fusion proteins between lysin and Sag1p would be fully translated, integrated into the Pp glycocalyx via the Sag1p part and thus displayed on the surface of the Pp cells. For the other 50%, translation would stop at the F2A sequence, resulting in a fully secreted LysM-CHAP protein. Selection of transformed Pp cells was achieved by complementation of histidine deficiency of the GS115 strain in histidine-free minimal medium by the HIS4-gene present on the pPIC9K backbone.

The library of recombined LysM-CHAP homologues was used to transform Pp cells, followed by selection of Pp clones expressing highly displayed LysM-CHAP sequences by fluorescence activated cell sorting (FACS) in the following way: Yeast library cultures were grown into stationary phase in 5-8 mL minimal medium (0.1 M phosphate buffer (pH 6.0), 1.34% YNB, 0.00004% biotin) in 50 mL falcon tubes sealed with a semi-permeable membrane at 28 °C and 250 rpm over night. The next day, cell densities were measured by absorbance. 4-8 OD_600_ units of cells were harvested by centrifugation (2000 xg 5min) and resuspended at a density of 0.5-1.0 in inducing medium (1.34% YNB, 0.1M NaPO_4_ pH 6.0, 0.00004% Biotin, 2% methanol and water). Cells were induced for approximately 16 - 22 h.

After induction, Pp cells were harvested and stained for FACS. Cells were collected by centrifugation (2000 xg, 5 min) and washed three times with 1 mL of freshly prepared cold sterile FACS buffer (1x DPBS, 1% BSA and 0.5mM EDTA). After the third wash, cells were adjusted to an OD_600nm_ of 1.0 in FACS buffer and stained with 1 µg/mL anti-V5-Alexa647 (MCA1360A647, Bio-Rad) for 1 hour while rotating protected from light at 4°C. Cells were subsequently washed three times with FACS buffer, resuspended at OD_600nm_ 1.0 to 2.0 and strained through a 0.40 µm mesh filter to reduce clumping.

Yeast cells were sorted on a BD FACSAria III (Becton Dickinson, Heidelberg, Germany) or a BD FACSMelody cell sorter with support from the MPL Core Facility (BioOptics – FACS, Max Perutz Labs, Vienna, Austria). A non-expressing gate was set after recording the background signals of a stained cell suspension transformed with the empty yeast display vector.

The top 0.5% of LysM-CHAP displaying Pp clones were sorted, followed by dilution and plating on yeast plates that had been supplemented with a final OD_600nm_ of 4.0 of autoclaved Sa ATCC 43300 and containing 0.1 M phosphate buffer (pH 6.0), 1.34% YNB, 0.00004% Biotin and 1% methanol for induction and 1.5% agar. Pp cells were diluted prior to plating to achieve a density per plate of around 150 colonies. Plates were incubated at 28 °C for 5-7 days, after which clearing zones (i.e. halos) formed around some Pp colonies. Formation of a halo around a Pp colony was due to the secretion of active LysM-CHAP, which cleaves the turbid Sa peptidoglycan embedded in the surrounding agar, leaving behind a translucent clearing zone, with broader clearing zones corresponding to higher LysM-CHAP secretion and activity.

100 colonies with halos were picked and sequenced by Sanger sequencing (Sort 1 sequencing). They were further pooled at equal density, induced to express lysins and subjected to a second round of FACS and plating on agar plates supplemented with autoclaved Sa, as described above. 100 colonies with halos were picked and sequenced (Sort 2 sequencing). Each unique sequence in the Sort 1 and Sort 2 sequencing data sets was counted. The counts were used to determine the enrichment of each sequence between Sort 1 and Sort 2. The top 10 enriched sequences were then used as the input for DNA shuffling for the next round of directed evolution. In this case, the input DNA of each selected variant was adjusted in proportion to its frequency observed in the Sort 2 sequencing dataset.

A total of two rounds of directed evolution was performed in this way. The top hits from each round were codon optimized for expression in human cells, and their HEK secretion and activity tested after expression in Expi293 cells (as described below), leading to the selection of the most secreted and active variants L1 and L2 from the first and second round of directed evolution, respectively.

### Construct design

Constructs intended for cytosolic production in Escherichia coli (Ec) were cloned into the pet29b vector (Novagen) under control of the IPTG-inducible T7 promoter. Amino acid sequences of the lysins were appended with an N-terminal hexa-histidine purification tag (N-HHHHHH-C) and a precision protease cleavage site (N-SAGLEVLFQGP-C) and codon optimized for expression in Ec.

Constructs for functional screening and expression in human embryonic kidney (HEK) cell culture were generated in the following way: DNA sequences encoding the lysin amino acid sequences were codon-optimized for human expression and cloned under control of the constitutive CMV promoter into pCDNA3.4 (Thermo Fisher Scientific). The constructs were appended with a 5’ Kozak sequence, followed by the coding sequence of a signal peptide derived from murine Ig kappa light chain precursor protein (mutant A2; sequence: N-MDMRAPAGIFGFLLVLFPGYRS-C)^5^, a “TG” dipeptide to enhance cleavage of the signal peptide after entering the secretory pathway, a hexa-histidine purification tag (N-HHHHHH-C) and a precision protease cleavage site (N-SAGLEVLFQGP-C).

In the case of lysins tagged with an Fc-domain, the protease cleavage site was followed by the coding sequence corresponding to human IgG1 Fc domain (amino acids 218-449 relative to UniProt-ID: P0DOX5) containing the Cys 222 Ser mutation and different linker sequences, followed by the first amino acid of the LysM-CHAP sequence.

For constructs containing a monomeric Fc-domain, the human IgG1 Fc domain lacking the hinge (amino acids 233-449 relative to UniProt-ID: P0DOX5) and containing the following mutations was used: Leu353Tyr, Thr368Tyr, Leu370Ala, Pro397Arg, Phe407Arg, Tyr409Met and Lys411Ala^6^.

For the production of heterodimeric Fc-hel8-L1-3 derivatives, the N-terminal hexa-histidine tag was substituted by a strep-tag (N-WSHPQFEK-C). Such constructs were designed in presence or absence of SpA inactivating RF-mutations^7^. To generate single arm heterodimers, Fc-hel8 constructs with an N-terminal strep-tag (with or without RF-mutations) and lacking the LysM-CHAP lysin part were generated.

Constructs intended for use in mRNA production were cloned into a proprietary template vector containing a 5’ T7-polymerase promoter, a 3’ polyA tail and restriction sites for template linearization. Constructs used for mRNA production were identical in amino and DNA sequence relative to their pCDNA3.4 derivatives but lacked the sequence corresponding to the cleavable purification tag.

### Protein purification and quality control

Purification of lysins for crystallography. Constructs encoding lysin CHAP domains (L1, L2) or a LysM-CHAP fusion (L1-3) and an N-terminal 3C-cleavable His6 affinity tag for purification were used to transform Ec BL21 (DE3) (NEB C2527H) competent cells, following the manufacturer’s protocol, then overexpressed and purified from cell lysate supernatant, following cell lysis via sonication. Briefly, transformed cells were grown in LB-carbenicillin to an optical density at 600 nm (OD) of 0.6-0.8, then 1 mM IPTG was added to induce expression at 18°C for 18 h. Cells were harvested via centrifugation and cell pellets were frozen and stored at -80°C. Frozen cell pellets were resuspended and thawed in lysis buffer (20 mM HEPES pH 7.4, 1 mM magnesium chloride, 20 mM imidazole), with 1 mM PMSF and benzonase added. Following sonication, cell lysate supernatant was separated and applied to an 0.45 µm filter membrane.

N-terminally His6-tagged lysin constructs were initially purified via immobilized metal affinity chromatography (IMAC) using a batch binding method. Briefly, 20 mL of 50% (v/v) slurry of Ni-NTA agarose beads were equilibrated with IMAC equilibration buffer (25 mM Tris-HCl pH 8.0, 300 mM NaCl, 10% glycerol). The equilibrated beads were added directly to the filtered lysate and allowed to shake gently at 4°C for 2 h. The beads were then washed with IMAC wash buffer (25 mM Tris-HCl pH 8.0, 300 mM NaCl, 10% glycerol, 15 mM imidazole), then transferred to a 20 mL disposable column and allowed to drain via gravity flow to pack the beads. After draining, 5 column volumes (CV) of IMAC wash buffer 2 (25 mM Tris-HCl pH 8.0, 300 mM NaCl, 10% glycerol, 30 mM imidazole) were added and allowed to drain completely via gravity flow. Protein was eluted by adding 3 CV of IMAC elution buffer (25 mM Tris-HCl pH 8.0, 300 mM NaCl, 10% glycerol, 250 mM imidazole). Presence of target protein was determined by SDS-PAGE analysis and A280 measurement of the elution pool.

Pooled protein was dialyzed against 1 L of HBS (20 mM HEPES pH 7.4, 150 mM NaCl) via a 10 kDa MWCO dialysis cassette at 4°C for 1 h. After 1 h, a 1:100 (3C:target protein) (w/w) ratio of HRV-3C protease was added and the cassette was dialyzed against 2 L of HBS buffer at 4°C for 18 h to ensure complete cleavage. Cleaved lysin constructs were further purified via size exclusion chromatography using a Superdex 75 16/600 column (equilibrated in HBS). Fractions of the major size exclusion chromatography peak, corresponding to the tagless lysin constructs, were pooled, concentrated, and verified for purity via SDS-PAGE analysis. Final purified samples were stored at 4°C until use for crystallization screening.

Protein purification for biophysical and functional characterization and for in vivo studies. The following proteins were purified from Expi293 supernatants: L1, L1-1, L1-2, L1-3, L2-1, Fc-hel8-L1, Fc-hel8-L1-1, Fc-hel8-L1-2, Fc-hel8-L1-3, Fc-hel8-L2-1, Fc-hel8-L2-2, Fc-hel8-L2-3, Fc-hel8-L2-4, all variants shown in Figure 1e, Fc-hel14-L1-5, Fc-hel14-L1-6 and PGLALA derivatives of Fc-hel8-L1-3 and Fc-hel14-L1-6. 30-800 mL cultures of Expi293 cells (Thermo Fisher Scientific) were grown shaking (120rpm) at 37 °C and 5 % CO_2_ in Expi293 medium according to the manufacturer’s instructions. Transfections were performed using the Expifectamine transfection kit (Thermo Fisher Scientific, A14524) and Optiplex (Thermo Fisher Scientific, A4096801) according to the manufacturer’s instructions and at a final DNA concentration of 1 µg/mL. Enhancer was added after 24 h and supernatants were harvested 72 h post transfection by centrifugation (500 x g, 30 min). Supernatants were clarified by a second highlZIspeed centrifugation (15000 x g, 30 min, 4°C). Cleared supernatants were adjusted to IMAC loading conditions by supplementation with NaCl (300 mM final), imidazole (20 mM final), and reducing agent (1 mM final). Benzonase was added (Merck Millipore, 70746-3, 1:10,000, v/v) to minimize nucleic acid contamination prior to chromatography. Proteins were purified directly from clarified culture supernatants using Ni²⁺lZIcharged HisTrap HP columns (5 mL; Cytiva) on an ÄKTA pure chromatography system. Columns were equilibrated in binding buffer (20 mM HEPES pH 7.0, 300 mM NaCl, 20 mM imidazole, 1 mM dithiothreitol (DTT)). Samples were loaded at 1.5 mL/min, followed by washing the column with binding buffer until baseline absorbance was reached. Bound proteins were eluted using step gradient changes to 20% and 80% of elution buffer (20mM HEPES pH 7.0, 300 mM NaCl, 20 mM imidazole, 1 mM DTT). Elution fractions were analyzed by SDS–PAGE followed by Coomassie staining and fractions containing the target protein were pooled. Pooled eluates were supplemented with fresh 1 mM DTT and HRV 3C protease (produced in-house) at a 1:100 protease:substrate molar ratio, followed by dialysis against formulation buffer (20 mM HEPES pH 7.0, 150 mM NaCl) at 4 °C against 300 sample volumes. Samples were finally sterilelZIfiltered (0.2 µm cutoff filters), aliquoted and frozen at -80 °C for long-term storage. Protein concentrations were determined by UV absorbance using molar extinction coefficients calculated from the amino acid sequence of each lysin variant using Geneious software.

Protein purification for exebacase and lysostaphin. Plasmids encoding lysostaphin or exebacase were transformed into chemically competent Ec BL21(DE3) or LEMO21(DE3) cells (New England BioLabs) by heat shock. Cells were incubated with plasmid DNA on ice for 301min, heatlZIshocked at 421°C for 101s, recovered in SOC or LB medium for 11h at 371°C, and plated on LB agar containing kanamycin (501µg1ml⁻¹) or kanamycin plus chloramphenicol (301µg1ml⁻¹) as appropriate. Protein expression was performed using autolZIinduction medium based on modified ZYMlZI5052. Starter cultures were grown in LB with antibiotics at 37 °C and diluted 1:200 into autolZIinduction medium supplemented with glycerol, glucose, and lactose. Expression cultures (2 L total) were incubated at 251°C for 241h with shaking. Cells were harvested by centrifugation at 4 °C. Cell pellets were resuspended in lysis buffer (201mM HEPES pH17.0, 3001mM NaCl, 201mM imidazole) and lysed by sonication. Lysates were treated with Benzonase, clarified by centrifugation, and applied to a 5 ml HisTrap HP column on an ÄKTA pure system. After washing with 40 mM imidazole, bound proteins were eluted using a stepwise imidazole gradient (200 and 800 mM). Fractions containing the target protein were pooled and analyzed by SDS–PAGE.

Purification of isoforms of Fc-hel8-L1-3 using strep tag affinity purification. Strep1tagged lysins were purified by affinity chromatography using StrepTrap XT 1 mL columns (Cytiva) on an ÄKTA pure 25 system. Columns were equilibrated in binding buffer (Buffer A), samples were loaded at 0.5 mL min⁻¹, washed with 10 CV of Buffer A, and eluted with Buffer A supplemented with 50 mM desthiobiotin (Buffer B). Buffer A consisted of 20 mM HEPES pH 7.0, 300 mM NaCl, and 1 mM DTT; Eluted proteins were pooled, concentrated as needed, and subjected to downstream processing and quality control.

Size exclusion chromatography. Purified proteins were analyzed for mono-dispersity by sizelZIexclusion chromatography on an ÄKTA pure 25 system (Cytiva) using a Superdex 200 Increase 10/300 GL column. The column was equilibrated in 20 mM HEPES (pH17.0) and 1501mM NaCl. Protein samples were clarified prior to injection by centrifugation and eluted isocratically at 0.5 mL/min. Elution was monitored by UV absorbance at 280 nm, and fractions containing the target protein were pooled for further analysis.

Nano Differential Scanning Fluorimetry (NanoDSF). Thermal stability of purified proteins was assessed using a Prometheus NT.48 instrument (NanoTemper Technologies). Protein samples were prepared at a final concentration of 0.5 mg/mL in formulation buffer. Samples were thawed immediately prior to measurement. Standard capillaries (NanoTemper Technologies) were filled with approximately 10 µL of sample and loaded into the instrument. All measurements were performed in duplicates. Samples were heated from 20 °C to 95 °C at a constant ramp rate of 1.0 °C/min. Intrinsic protein fluorescence was monitored with excitation at 280 nm and emission recorded at 330 and 350 nm. Thermal unfolding transitions were determined from changes in the fluorescence emission ratio (F₃₅₀/F₃₃₀). In parallel, changes in light scattering were monitored as attenuation units to assess sample aggregation or turbidity during thermal unfolding.

## Protein quality control

SDS–PAGE purity assessment. Lysin purity and integrity were evaluated by SDS–PAGE under reducing and non1reducing conditions. Samples (2.5 µg) were resolved on AnykD Criterion TGX precast gel (BioRad 5671125) and visualized by InstantBlue® Coomassie Protein Stain (Abcam ab119211) staining. Purity was estimated by densitometric analysis using ImageQuantTL software, and preparations exhibiting ≥95% purity were used for further analyses.

Size-exclusion chromatography (SEC-HPLC). Size distribution and aggregation were assessed by size1exclusion chromatography using a TSKgel® G3000SWXL column (5 µm) coupled to an UltiMate™ 3000 HPLC system. Proteins were eluted in 20 mM HEPES, 150 mM NaCl, 300 mM Arginine pH7 buffer at a flow rate of 0.3 mL/min, and elution profiles were monitored at 280 nm. Monomer content was determined by peak area integration and was required to exceed 95%.

Intact mass analysis. Protein identity was confirmed by intact mass analysis using high1resolution LC-MS. Samples were reduced in 3 M guanidine hydrochloride containing 12 mM dithiothreitol for 15 min at 65 °C, followed by dilution with water prior to analysis. Reduced proteins (2 µL) were separated on a Waters BioResolve Polyphenyl mAb column using a Vanquish Horizon UHPLC system operated at a flow rate of 250 µL/min with a step gradient of 15–55–80% acetonitrile. Mass spectra were acquired on a Synapt G21Si mass spectrometer in resolution mode. Average mass spectra were reconstructed using the MaxEnt1 algorithm and compared with theoretical masses to confirm protein identity.

CMK peptide labelling of the CHAP active site. The redox state of the catalytic cysteine within the CHAP domain was assessed by covalent labeling with a chloromethyl ketone (CMK) peptide probe. Purified lysins (4 µM) were incubated in protein buffer (20 mM HEPES, pH 7.0, 150 mM NaCl) with CMK peptide at a final concentration of 40 µM for 1 h at room temperature (RT) in the dark. Reactions were quenched by addition of reducing Laemmli sample buffer containing 21mercaptoethanol, followed by heating at 90 °C for 10 min. Labeled proteins were resolved by SDS–PAGE for qualitative assessment of CMK modification. As CMK selectively reacts with reduced, catalytically competent cysteine residues, labeling was used to evaluate the redox state and functional integrity of the cysteine-dependent CHAP catalytic core.

Endotoxin Assessment. Recombinant proteins used for in vivo studies were tested for endotoxin contamination using a Limulus amebocyte lysate (LAL) assay (Pierce Chromogenic Endotoxin Quantitation Kit, Thermo Scientific, A39552) and were confirmed to be suitable for in vivo use.

## Generation of LysM-CHAP reactive antibodies

Antigen Preparation. The C-terminal LysM-CHAP fragment of LytN was expressed in Ec and purified by Ni-NTA affinity chromatography as described above. Protein eluates were treated with HRV 3C protease (produced in-house, 1:100 protease:lysin molar ratio) while being dialyzed over night against 300 sample volumes of final storage buffer (20 mM HEPES pH 7.0, 150 mM NaCl). Purity (≥85%) was confirmed by SDS–PAGE followed by Coomassie staining. Aliquots were stored at –80 °C until immunization.

Rabbit Immunization. Polyclonal antibodies were raised by Genscript in New Zealand White rabbits under a 3lZIimmunization protocol conducted in an AAALAC/OLAWlZIaccredited facility. PrelZIimmune serum was collected before subcutaneous administration of antigen; two boosters were given at standard intervals; postlZIimmune serum was collected after the final boost.

Affinity Purification and Crossl/Adsorption. Pooled antisera (401mL per rabbit) were subjected to antigenlZIaffinity purification, yielding LytN–specific IgG in DPBS (pH17.4) with 0.02% ProClin1300 and ≥99% purity by SDS–PAGE. To remove unwanted antilZIIgG reactivity, antibodies were sequentially crosslZIadsorbed against immobilized human and mouse total IgG (≥85% purity). The final preparation yielded 11.441mg total IgG at 0.4871mg/mL.

Biotinylation. An aliquot (2 mg) of purified IgG was biotinlZIconjugated by incubation with a 10-fold molar excess of biotin-sulfo-NHS for 30 min at 23 °C, followed by buffer exchange into DPBS. The antibody solution (final Ig concentration at 0.368 mg/mL) was supplemented with 1% BSA and 0.02% ProClin 300, aliquoted and frozen at -80 °C until use.

## Synthesis of Staphylococcus aureus (Sa) stem peptide analogues

Pep1a was designed as a full-length Sa stem peptide appended with an N-terminal EDANS fluorophore and a C-terminal DABCYL quencher. The peptide was synthesized at Genscript at >95% purity with the following sequence code: {GLU(EDANS)}A{D-GAMMA-GLN}K{D-ALA}GGGGG{LYS(DABCYL)}. The lyophilizate was dissolved at 1 mM final concentration in DMSO. Aliquots were stored at -80 °C until use.

Pep1a-CMK was designed as a fragment of the Sa stem peptide containing positions 1-4 (Figure S3b) with an N-terminal 5(6)-FAM fluorophore and a C-terminal chloromethyl-ketone group. Peptide synthesis was performed at JPT peptide Technologies GmbH using the sequence code “Acetyl-{Lys(5/6FAM)}Ala{D-GAMMA-GLN}Lys{D-ALA}-Chloromethylketone”. The peptide was synthesized and purified in 10 mg scale at >70% target purity, aliquoted and lyophilized, followed by storage at -80 °C. Aliquots of the peptide were resuspended in 100% anhydrous DMSO at a final concentration of 1 mM. Working aliquots of this solution were stored at -80 °C until use.

## Peptidase assays using Pep1a

Peptidase activities of LysM-CHAP derivatives were monitored using Pep1a as a stem peptide substrate. The close proximity between the fluorophore and quencher in the peptide results in quenching of EDANS fluorescence. Cleavage of the peptide results in the release of the quencher from the fluorophore and thus in an increase in fluorescence. The change in fluorescence can be quantified over time to determine rates of peptide hydrolysis by different lysins.

2.5 µM purified lysin (L1-3 and L2-1) was mixed with increasing concentrations of Pep1a in assay buffer (final concentrations: 20 mM HEPES, pH 7.0, 52.5 mM NaCl, 1 mM CaCl2, 1 % bovine serum albumin and 5.5 % DMSO), followed by fluorescent measurement (excitation 340 nm, emission 460 nm) in a plate reader (Tecan) at 37 °C. The fluorescent signal of fully cleaved peptide was determined in a separate condition in which recombinant lysostaphin was incubated at 2.5 µM with corresponding substrate concentrations (data not shown). The signal plateau reached with lysostaphin was used to determine the correlation between fluorescent signal and substrate concentration. Rates of substrate hydrolysis for lysin variants were determined by fitting the initial linear phase of the fluorescent signal curves. The rates of signal changes were converted to rates of changes in substrate concentration and plotted against the input substrate concentration. These data was then fit to the Michaelis Menten model in Graphpad PRISM to estimate the Michaelis Menten constant K_M_.

## Small angle X-ray scattering of Fc-hel8-L1-3

Small-angle X-ray scattering (SAXS) data were acquired by BIOSAXS GmbH at the P12 beamline of the PETRA III synchrotron (DESY, Hamburg, Germany). The undulator (gap 20.95 mm) and monochromator were tuned to an incident energy of 10 keV. Beamline calibration of the angular axis was performed using silver behenate. Prior to data collection, beamline performance was verified following the P12 standard operating procedures, including measurements of empty capillaries, Milli-Q water, bovine serum albumin (BSA), and the corresponding HEPES buffer.

Technical Specifications of P12 beamline used for the analysis.

|  |  |
| --- | --- |
| Source | Petra III U29 undulator, P12 BioSAXS beamline |
| Detector | hybrid photon detector Pilatus 6M |
| Energy range | 10 keV |
| Maximum flux at sample | 1*10<sup>12</sup> photons per second |
| Beam size at sample | 250 μm × 100 μm |
| s-range | 0.03 – 7.25 nm<sup>-1</sup> |
| Wavelength | 0.124 nm |
| Sample-to-Detector Distance | 3 m |
| Number of bunches | 40 |
| PETRA III ring current | 100 mA |
| Axis Calibration Standard | silver behenate |
| Beamstop position | off-set |

Samples were subjected to inline size-exclusion chromatography–coupled SAXS (SEC–SAXS) using an Agilent Bioinert Infinity 1260 HPLC system connected to a Cytiva Superdex 200 Increase 5/150 column. The column was equilibrated with 10 CV of SEC buffer (20 mM HEPES, 150 mM NaCl, pH 7.0) and linked directly to the SAXS flow cell. For each SEC–SAXS run, 25 μL of protein sample at 5 mg/mL were injected. A total of 1,800 scattering frames were recorded during elution of the protein peak and subjected to automated radial averaging as previously described^8,9^. Data processing and extraction of global scattering parameters were performed using CHROMIXS^10^ and PRIMUS^11^, including calculation of real-space distance distribution functions. Ab initio shape reconstruction was carried out as described by Svergun et al, 1999^12^ (data not shown), yielding overall shapes of Fc-hel8-L1-3 with the following parameters:

|  |  |
| --- | --- |
| $R_{g1}$ (nm) | $6.4 \pm 0.1$ |
| $R_{g2}$ (nm) | $7.2 \pm 0.1$ |
| $D_{max}$ (nm) | $30 \pm 2$ |
| $V_{dam}$ (nm<sup>3</sup>) | $240 \pm 15$ |
| $MW_{dam}$ (kDa) | $120 \pm 10$ |

with R_g1_ being the radius of gyration determined from the Guinier approximation, R_g2_ the radius of gyration determined from the distance distribution function computed by the program GNOM^13^, D_max_ the maximum particle size assessed from the p(r) function in GNOM, V_dam_ the volume of a typical ab initio model reconstructed by the program DAMMIN^12^ and MW_dam_, the approximate molecular weight assessed from the typical DAMMIN model.

Rigid-body modelling of Fc-hel8-L1-3 was performed with CORAL^14^. Fc-hel8-L1-3 was represented as a dimer comprising the IgG1 Fc domain (PDB: 6eaq), a helical linker derived from PDB:6jfv, and the LysM and CHAP domains (PDB: 11CI), with the latter two connected by a flexible linker. The dimeric nature of Fc was placed at the origin and oriented with the principal axes along X, Y and Z. Multiple CORAL runs were conducted on the Fc-hel8-L1-3 data and the restored symmetric dimers displayed an overall similar appearance yielding a good fit to the data. Two different organizations of the monomers in the dimer were observed, namely trans-dimers (as shown in Figure 2b) and cis-dimers (data not shown), with the LysM-CHAP arms located either on the opposite or same site, respectively, of the fused Fc domain. Both isoforms displayed otherwise similar conformations and dimensions.

## X-ray crystallography

General X-ray crystallographic methods. Crystallographic datasets were collected at beamline P13 and P14 operated by EMBL Hamburg at the PETRA III storage ring (DESY, Hamburg, Germany) and beamline CMCF-ID operated by the University of Saskatchewan at the Canadian Light Source national research facility. Beamline CMCF-ID at the Canadian Light Source is supported by the Canada Foundation for Innovation (CFI), the Natural Sciences and Engineering Research Council (NSERC), the National Research Council (NRC), the Canadian Institutes of Health Research (CIHR), the Government of Saskatchewan, and the University of Saskatchewan. All diffraction images were processed using a xia2/DIALS pipeline^15,16^. Initial phases were obtained by molecular replacement using the PhaserMR program, found within the CCP4 software suite^17^. Model building and refinement were performed using Coot and Refmac5, respectively^18,19^. Final structures were validated for rotamer and Ramachandran outliers using the Molprobity server^20^ . Figures were prepared using the PyMOL molecular graphics software package^21^. Contact mapping of the LysM:CHAP interface was performed using NCONT (CCP4 suite) and PISA server^22^

Crystallization and structure solution of L2 CHAP domain. Initial crystallization screens of the L2 CHAP domain were conducted at a concentration of 10 mg/mL in storage buffer [20 mM HEPES pH 7.4, 150 mM NaCl, and 2 mM TCEP]. The sample was mixed in three ratios (1:2, 1:1, and 2:1) with five screening kits, containing 96 conditions each, to a final volume of 150 nL per drop using an SPT Labtech mosquito xtal3 robot. Crystal formation was observed using automated imaging over the course of 28 days. Thin plate crystals were observed via sitting drop vapor diffusion within three days in the Wizard 3/4 screening kit (Molecular Dimensions), in well A10 using a mixing ratio of 1:1 protein-to-well. Crystals were further optimized in large format sitting drop trays, with the best diffracting crystals appearing in the final crystallization condition [18% PEG 3350, 0.2 M potassium thiocyanate]. Crystals were then harvested from a brief soak in cryoprotectant solution (well solution supplemented with 15% (v/v) glycerol) using rayon loops and flash frozen in liquid nitrogen. Initial phases were obtained using a literature crystal structure of a LysK CHAP domain as the initial search model (PDB accession code 4CSH).

The L2 CHAP domain crystallizes in the C2221 space group with one copy in the asymmetric unit. The final model contains residues 74-215 (correlating with the residue numbering of the LysM-CHAP protein) in chain A, 1-100 in chain B (water molecules), and 1-12 in chain C (sodium ions and thiocyanate monomers). Residues 137-141 within chain A are disordered, lacking electron density, and not modelled in the final structure. Surface-facing residues without appreciable electron density for their side chains were stubbed at the α-carbon. Of the residues modelled, 100% are in allowed or preferred regions indicated by Ramachandran validation statistics.

Crystallization and structure solution of L1 CHAP domain. The purified L1 CHAP domain was concentrated to 27 mg/mL in storage buffer [20 mM HEPES pH 7.4, 150 mM NaCl, and 2 mM TCEP], for crystallization screening. After two days at high concentration at 4°C, crystals were observed in the stock tube of the concentrated sample. The presence of protein crystals was supported by the lowered concentration of protein within the soluble phase of the sample and observation of a hexagonal pyramid crystal morphology. Crystals were then harvested from a brief soak in cryoprotectant solution (storage buffer supplemented with 15% (v/v) glycerol) using rayon loops and flash frozen in LN2. Initial phases were obtained using an internal crystal structure of the L2 variant of the lysin CHAP domain (PDB accession code 11CG).

The L1 CHAP domain crystallizes in the P1 space group with four copies in the asymmetric unit. The final model contains residues 76-215 (correlating with the residue numbering of the LysM-CHAP protein) in chain A, 74-215 in chain B, 76-214 in chain C, 74-215 in chain D, 1-645 in chain E (water molecules), 1-13 in chain F (sodium ions), and 1-6 in chain G (glycerol molecules). Surface-facing residues without appreciable electron density for their side chains were stubbed at the α -carbon. Of the residues modelled, 100% are in allowed or preferred regions indicated by Ramachandran validation statistics.

Co-Crystallization and structure solution of L1 CHAP:CMK complex. Co-crystallization was performed by generating a 2:1 ligand-to-protein mixture of the purified L1 CHAP domain and Pep1a-CMK ligand at a concentration of 2.5 mg/mL. After a 15-minute incubation period on ice, the sample was run on a PD-10 desalting column to remove unbound ligand, eluted with storage buffer [20 mM HEPES pH 7.4, 150 mM NaCl, 2 mM TCEP], and concentrated to 30 mg/mL to be used for crystal screening. The sample was mixed in three ratios (1:2, 1:1, and 2:1) with four screening kits, containing 96 conditions each, to a final volume of 150 nL per drop using an SPT Labtech mosquito xtal3 robot. Crystal formation was observed using automated imaging over the course of 28 days. Bright orange cubic crystals were observed via sitting drop vapor diffusion within one day in the ProComplex Suite screening kit (NeXtal), in well H1, and was present in all three mixing ratios. Crystals were further optimized, with the best diffracting crystals appearing in the final crystallization condition [2.4 M NaCl, 0.1 M sodium citrate pH 6.0]. Crystals were then harvested from a brief soak in cryoprotectant solution (well solution supplemented with 15% (v/v) glycerol) using rayon loops and flash frozen in LN2. Initial phases were obtained using an internal crystal structure of the apo L1 CHAP domain (PDB accession code 11CF).

The L1 CHAP:Pep1a-CMK covalent complex co-crystallizes in the P42212 space group with one copy in the asymmetric unit. The final model contains residues 74-215 (correlating with the residue numbering of the LysM-CHAP protein) and residue 270 (full Pep1a-CMK ligand) in chain A, 1-7 in chain B (sodium and thiocyanate ions), and 1-131 in chain C (water molecules). Surface-facing residues without appreciable electron density for their side chains were stubbed at the α-carbon. Of the residues modelled, 100% are in allowed or preferred regions indicated by Ramachandran validation statistics. Electron density for the Pep1a-CMK ligand (residue 270, chain A) is absent from C16 to C26, which includes the N-acetyl group, lysine linker, and first carbon ring (lactone ring) of the FAM fluorophore (Figure 3c). Weak electron density was present for the tricyclic xanthene backbone of the fluorophore. The portion of the ligand lacking density was modelled in most likely conformation using real space refinement in Coot, given crystal packing constraints.

Co-Crystallization and structure solution of L1-3:CMK complex. Co-crystallization was performed by generating a 2:1 ligand-to-protein mixture of the purified L1-3 protein and Pep1a-CMK ligand at a concentration of 1 mg/mL. After a 15-minute incubation period on ice, the sample was run on a PD-10 desalting column to remove unbound ligand, eluted with storage buffer [20 mM HEPES pH 7.4, 150 mM NaCl, 2 mM TCEP], and concentrated to 6.5 mg/mL to be used for crystal screening. The sample was mixed in three ratios (1:2, 1:1, and 2:1) with five screening kits, containing 96 conditions each, to a final volume of 150 nL per drop using an SPT Labtech mosquito xtal3 robot. Crystal formation was observed using automated imaging over the course of 28 days. Faint orange plate-clustered crystals were observed via sitting drop vapor diffusion within 2 days in the PEGS II Suite screening kit (NeXtal), in well G1 using a mixing ratio of 1:2 protein-to-well, in the final crystallization condition [25% PEG MME 5000, 0.1 M TRIS pH 8.5, and 0.2 M lithium sulfate]. Crystals were then harvested from a brief soak in cryoprotectant solution (well solution supplemented with 15% (v/v) glycerol) using rayon loops and flash frozen in LN2. Initial phases were obtained using an internal crystal structure of the L1 CHAP domain co-crystallized with the Pep1a-CMK ligand (PDB accession code 11CH) and an AlphaFold2 model of the LysM domain encoded in L1-3.

The L1-3:Pep1a-CMK covalent complex crystallizes in the C2221 space group with one full complex in the asymmetric unit. The final model contains three chains (A, B, and C). Chain A contains residues 6-52 (belonging to the L1-3 LysM domain), residues 74-215 (belonging to the L1-3 CHAP domain), and residue 270 (Pep1a-CMK). Residues 53-73 of chain A, which comprise the linker between domains, are disordered and absent from the modelled structure. Chain B contains four water molecules and chain C contains one sodium ion. Surface-facing residues without appreciable electron density for their side chains were stubbed at the α--carbon. Of the residues modelled, 100% are in allowed or preferred regions indicated by Ramachandran validation statistics. Strong electron density for the Pep1a-CMK ligand (residue 270, chain A) ends at carbon 18, just before the lysine linker to the FAM fluorophore. The portion of the ligand lacking density was modelled in most likely conformation using real space refinement in Coot, given crystal packing constraints, but exhibits a large degree of flexibility as intended by the design of the inhibitor/fluorophore.

## Measurement of cell binding kinetics by biolayer interferometry (BLI)

BLI experiments were performed using an eight-channel Octet® R8 instrument (Sartorius). All measurements were conducted at 30 °C in assay buffer containing 20 mM HEPES (pH17.0), 1501mM NaCl, 0.0025% (v/v) Tween-20, and 0.075% (w/v) BSA. Samples were formatted in a 96-well black plate (Greiner Bio-One), with each well containing 200 µL. For each experiment, eight streptavidin (SA) biosensor tips (Sartorius) were used. Sensors were first equilibrated in assay buffer for 10 min and then loaded with biotinylated Sa ATCC 43300 cells prepared at a 1:16 dilution from a stock with an OD of 0.8 in 25% glycerol. After ligand loading, sensors were quenched in biocytin (5 µg/mL) to block residual streptavidin sites and minimize nonspecific binding. Sensors were then dipped into analyte solutions containing catalytically inactive endolysins at varying concentrations (1.56–100 nM) prepared in assay buffer. Association and dissociation steps were performed for 120 s each. All experiments included reference measurements using unloaded SA biosensors dipped into the same analyte-containing wells, as well as zero-analyte controls for baseline correction. Data was processed and analyzed using Octet Analysis Studio software (version 13.0). Prior to fitting, datasets were reference-subtracted, baseline-aligned, and corrected for interstep shifts. Binding kinetics were determined using a 1:1 binding model with global fitting applied to the association and dissociation phases. The first 120 s of association and 120 s of dissociation were used for fitting. Rate constants (k_on_, k_off_) and equilibrium dissociation constants (K_D_) were calculated from the fitted curves. Responses values below 0.05 nm were taken to indicate no binding and sensograms exhibiting abnormal baselines were excluded from analysis.

## Active site engineering library design

For the Combinatorial Motif Engraftment (CME) approach of active site diversification, we first selected amino acids positions in the vicinity of the catalytic amino acids (Cys 98, His 161 and Asp 181), the substrate, the Ca^2+^ binding site and the P1 site (positions 93, 94, 96, 99, 117, 118, 122, 159, 162, 163, 181, 189, 196, 198). We then aligned Alphafold 2.0^23^ generated models of 62 diverse CHAP homologues via their catalytic residues and extracted the corresponding amino acids at the selected positions. We finally rationally defined functional motifs as sets of interacting residues. These homologue residue sets were then enumerated as non-redundant pairs, triplets, quadruplets and quintuplets and grafted onto the L1-3 CHAP domain, resulting in a library of 225 CHAP designs.

For the inverse folding approach using the DeepChain platform (InstaDeep Ltd., London, UK; https://deepchain.bio), we employed large scale in silico sequence diversification of the CHAP catalytic pocket, followed by multidimensional ranking using a set of optimized filters. Positions within 6 Å distance of the stem peptide were diversified by inverse protein folding into residues found in CHAP homologues using LigandMPNN^25^, and proteinDPO^26^ with the L1-3 crystal structure as a template. A total of ∼55 million CHAP sequences containing up to 12 simultaneous mutations were generated. The sequences were first filtered for absence of defined sequence elements (N-glycosylation sites, cysteines, prolines, protease cleavage sites, poly-glycine stretches) and then further scored based on LigandMPNN and ProteinDPO-sequence scores, the adherence to geometric constraints in the catalytic site (Cys 98-His 161 distance <=3.8 Å; His 161 – position 181 distance <= 3.4 Å, position 181 pKa <=-1; position 184 either Trp, Tyr, Phe, or His) and an activity predictor derived from training data of CHAP derivatives active against Sa, resulting in the selection of a total of 225 sequences for screening.

## High throughput screening procedure

DNA plating, transfection and harvest. Transfection plates were prepared and DNA concentrations normalized to 100 ng/µL using an epMotion 5073 automated liquid handling system. A total of 1µg DNA was mixed with transfection reagents in 96 well format, followed by transfer onto 1 mL Expi293 cultures in 24 well format. Transfections, enhancer addition and cell culture conditions were as described above. Supernatants were finally harvested and aliquoted into multiple 96 well plates. Plates were either processed directly or frozen at - 80 °C.

Quantification of lysin secretion levels by BLI. Lysins secreted into Expi293 supernatants were quantified via their Fc domains by BLI using an OctetR8 instrument (Sartorius) and AHC2 anti-human Fc sensor tips (Sartorius, 18-5143). The following buffers were prepared freshly on the day of the experiment: assay buffer (25% supernatant from mock-transfected Expi293 cells, 74.45% (v/v) 1x DPBS, 0.05% (v/v) Tween 20, 0.5% (w/v) BSA); dilution buffer (99.263% (v/v) 1x DPBS, 0.067% (v/v) Tween 20, 0.67 % (w/v) BSA); regeneration buffer (10 mM glycine pH 1.7, 0.05% (v/v) Tween 20). Prior to measurement, biosensors were equilibrated in assay buffer for 10 min at 30 °C. A standard curve was generated by preparing recombinantly purified Fc-hel14-L1-4 at two-fold serial dilutions between 200 – 3.125 µg/mL in assay buffer. Supernatant samples were diluted four-fold diluted in dilution buffer. A total of 200 µl of each sample was used for quantifications. Biosensor movement was programmed in the following way: 1.) regeneration of the sensor in regeneration buffer (5 s, 400 rpm). 2.) sensor wash in assay buffer (5 s, 400 rpm). 3.) steps 1 and 2 were repeated three times. 4.) sample measurement (120s – 400 rpm). Data was processed in Octet Analysis Studio software (version 13.0), which was used to fit k_on_ of binding between target protein and sensors. k_on_derived from recombinant standard recorded at different concentrations was used to determine a standard-curve by non-linear regression. The standard curve was then used to estimate the concentrations of lysins in the supernatants. Molecular weights of lysin variants were calculated from their amino acid composition (of the monomeric construct) and used to convert into molar concentrations.

High throughput growth inhibition assay. Overnight cultures of Sa ATCC 43300 and BAA-1717 were grown in tryptic soy broth (TSB) at 37 °C and shaking at 220 rpm, diluted 1:20 into fresh TSB to an optical density at 600 nm (OD) of ∼0.05, and further grown until reaching mid-log phase (OD ∼1.0). This culture was then diluted in cation-adjusted Mueller–Hinton broth supplemented with 20% heat-inactivated horse serum (caMHB + 20% HS) to ∼5 × 10^5^ colony forming units /mL (CFU/mL). 80 µL of supernatants were pre-dispensed into sterile flat-bottom 96-well plates. Using a TECAN Freedom EVO150 liquid handling platform equipped with a 96-channel head, 40 μL of caMHB + 20% HS was dispensed into each well of additional 96-well plates and a series of 2-fold serial dilutions of each supernatant sample was performed across these plates (the first plate contained undiluted samples). Subsequently, 60 μL of the prepared bacterial suspension was added to each well and plate, yielding a final assay volume of 100 μL per well (60% inoculum, 40% sample). Plates were covered and incubated at 37 °C for 18-19 h. Bacterial growth was then assessed by measuring OD in each well using a microplate reader (with 90 s of orbital shaking before reading). The minimal inhibitory concentration (MIC) of each sample was defined as the lowest sample concentration that completely prevented bacterial growth as judged by OD readings equivalent to sterile control wells.

Lysin purification from screening supernatants. Ala mutation (11–18,28,39,40–46) on LysM with FcZIhel14ZIL1ZI4 background in supernatants were purified using a membrane-based 24-well immobilized metal affinity chromatography plate (Capturem™ His-Tagged Purification 24-well Plate, Takara Bio, Cat. No. 635730) operated on a vacuum manifold. Cell culture supernatants containing His-tagged targets were supplemented with imidazole (201mM final) and Benzonase nuclease (1:10,000 dilution) and mixed with 1/10 volume of a 10× binding buffer (2001mM HEPES pH17.0, 31M NaCl, 2001mM imidazole) before clarification by centrifugation (500 × g, 101min). The Capturem plate was equilibrated by three successive vacuum filtrations of 21mL Buffer A per well (e.g., 201mM HEPES pH17.0, 3001mM NaCl, 201mM imidazole, 21mM β-mercaptoethanol). Clarified samples (0.5–4.51mL per well) were then loaded onto the plate and incubated for ∼301min to allow His-tagged proteins to bind to the membrane (with up to three re-applications of the flowthrough to maximize binding), followed by vacuum filtration to collect unbound material. Each well was washed at least twice with 21mL of Wash Buffer (95% Buffer A/5% Buffer B, where Buffer B is the same as Buffer A containing 11M imidazole) to remove non-specifically bound proteins. Bound proteins were eluted by adding 500 μL of Elution Buffer (50% Buffer A/50% Buffer B, yielding ∼5001mM imidazole) to each well, sealing the plate with an adhesive film, incubating for 20–301min at RT, and then applying vacuum to collect the eluates. The eluted proteins (∼0.81mL per well) were buffer-exchanged into final storage buffer (201mM HEPES, 1501mM NaCl, pH17.0, 11mM TCEP) using a 96-well size-exclusion filtration plate (Centripure 96, emp Biotech CP-0582), and the resulting samples were sterile-filtered through a 0.22 μm membrane (AcroPrep 24-well filter plate, VWR, 738-0221) by centrifugation at 4,000 × g for 21min. Finally, purified proteins were aliquoted into sterile 96-well polypropylene plates and stored at –801°C. Protein concentrations were measured using a NanoDrop spectrophotometer, and sample purity and yield were assessed by SDS–PAGE.

OD reduction assays. Sa BAA-1717 and ATCC 43300 were cultured aerobically on blood agar (BD) or in tryptic soy broth (TSB) medium at 37 °C. Pre-cultures were prepared by inoculating TSB medium with a few bacterial colonies grown on an agar plate, followed by overnight incubation. The pre-culture was used to inoculate 100 mL of TSB medium at an initial OD of 0.05. Cells were grown in baffled flasks at 37°C and 130 rpm until an OD of ∼1.1 was reached (requiring ∼ 2.5–3 h). The bacterial cells were subsequently harvested by centrifugation (4000 × g, 10 min). Cells were washed twice in DPBS supplemented 1 mM CaCl_2_ and 0.5 mM MgCl_2_ (Thermo Fisher Scientific, 14040), followed by resuspension in the same buffer at a final OD of 2. Supernatants were diluted 1:4 to 1:8 (depending on the expression levels) in Expi293 expression medium supplemented with 0.5 mM TCEP. 100 µL of the diluted supernatants were transferred into a flat-bottomed 96-well plate and mixed with an equal volume of the bacterial suspension, yielding a final OD of 1. The reaction plate was immediately placed into a Spectramax Plus plate reader (Molecular Devices) pre-heated to 37 °C. OD was then measured at intervals of 20 s over a period of 2 h. Rates of OD reduction were extracted as the maximum slope of the OD reduction curves. To mitigate the impact of noise in the OD reduction data, which can hinder accurate determination of the maximum slope, the data was smoothed using a cubic spline representation. The maximum slope was identified as the minimum of the first derivative of the spline. Data smoothing and slope determination were implemented in Python using the SciPy package to compute smoothing cubic B-splines. OD reduction rates were further normalized relative to a chosen lysin reference. For this purpose, rates of the reference construct were determined at different concentrations on each 96 well screening plate. A set of different regression models was fit to the concentration-dependence of the OD reduction rate in Python. The model with the highest confidence was selected. The rate of each construct was then normalized by the rate predicted for the reference at the concentration of the respective construct. This normalization yields a dimensionless factor, representing the activity of the construct relative to the reference.

Serum stability testing. 6 µL of Expi293 supernatants expressing linker 2 variants of Fc-hel8-L1-3 were mixed with either 54 µL of DPBS or pooled human serum (Dunne, ISER500 mL) in a 96 well PCR plate. The plate was sealed with translucent foil, placed in an anaerobic box together with an anaerobic atmosphere generation bag (Fisher Scientific, AN0025A) and incubated at 37 °C for 24 h. A 1 mM stock of Pep1a-CMK in DMSO was diluted to 1 µM in DPBS. 10 µL of the serum- or DPBS-treated supernatants were mixed with 50 µL of the Pep1a-CMK stock and incubated at RT for 10 min in the dark. The reaction was stopped by addition of SDS-PAGE sample buffer and analyzed by SDS-PAGE followed by fluorimetry using an Amersham Gel Imager with filter settings for fluorescence detection in the FAM channel.

## Bacterial strains and culture conditions

Staphylococci were streaked for single colonies from cryo vials onto Columbia Agar with 5% sheep blood (BD) and statically incubated overnight (o/n) at 37 °C. Plates were kept at 4 °C for up to 2 weeks. Multiple colonies were used to inoculate 10 mL TSB (Carl Roth) cultures in 50 mL Falcon tubes and incubated o/n at 37 °C with shaking at 220 rpm. o/n cultures were diluted in TSB to OD_600_=0.05 and incubated at 37 °C with shaking at 240 rpm for 2-3 h. Cells were harvested at OD=0.7-1.5 by centrifugation at 4000 g for 5 min and washed twice with DPBS supplemented 1 mM CaCl_2_ and 0.5 mM MgCl_2_ (Thermo Fisher Scientific, 14040), and resuspended as needed. The following strains were used for this study: Sa ATCC 43300 (ATCC), BAA-1717 (ATCC), NCTC 8178 (NCTC), S. epidermidis DSM3269 (DSMZ), S. lugdunensis DSM4804 (DSMZ), S. warneri DSM20316 (DSMZ) and S. capitis (clinical isolate).

## Growth inhibition assays for determination of minimal inhibitory concentrations (MICs)

### MIC was determined using a microbroth dilution method following the CLSI Standard M100-Ed35

Appendix H2 for antimicrobial susceptibility testing^27^. 2-fold dilution series of lysins were prepared as 5× stock solutions in cation adjusted MHB (caMHB, Sigma Aldrich) supplemented with 20% horse serum (heat-inactivated, Thermo Fisher Scientific, 26050070). Fc-tagged lysins were tested at 0.25-32 µg/mL (1-128 µg/mL for Fc-hel8-L1-3), naked lysins were tested at 0.125-16 µg/mL. Washed mid-log bacterial cells were adjusted to OD_600_=0.001 in caMHB with 20% horse serum, to reach a final inoculum of 5×10^5^ CFU/mL. Reactions were set up in triplicate in 96-well U-bottom plates by mixing 20 µL of lysin solution with 80 µL bacterial suspension. Plates were statically incubated at 37 °C for 17-19 h. MIC was defined as the lowest concentration of lysin that still prevented growth by visual inspection and determined for each replicate.

## Time-kill assay and determination of minimal eradication concentration (MEC)

Lytic activity of lysins against Sa alone or in combination with antibiotics (vancomycin, (MIP Pharma), cefazolin (Sigma Aldrich), and linezolide (Thermo Fisher Scientific)) was assessed by TKAs. Washed mid-log cells (see above) were resuspended to OD_600_=0.125 in DPBS supplemented with 1 mM CaCl_2_ and 1 mM MgCl_2_ (Thermo Fisher Scientific, 14040) or HuS (80% in the reaction; Innovative Research). This corresponded to approximately 5×10^7^ to 2×10^8^ CFU/mL in the reaction. Lysins (± antibiotic) were prepared as 5× stock solutions in lysin storage buffer with 2.5% BSA (resulting in 0.5% BSA in the reaction; Carl Roth). If more than one antibiotic was used, these were first mixed in appropriate concentrations in lysin storage buffer with 2.5% BSA to again obtain a 5x stock solution in combination with the lysin. Reactions were set up in duplicate in 96-well flat-bottom plates by mixing 20 µL lysin (± antibiotic) with 80 µL bacterial suspension. Plates were incubated statically in a tightly sealed plastic box with an anaerobic atmosphere generation bag (Fisher Scientific, AN0025A) and an anaerobic indicator (Thermo Fisher Scientific, BR0055B) at 37 °C for 3 h unless stated otherwise. Quantitative spotting was performed by five 10-fold serial dilutions in DPBS and spotting 2 µL of the dilutions onto TSA plates (BD). After o/n incubation at 37 °C, CFU/mL were determined. The log10 reduction in CFU/mL was calculated relative to the buffer-treated growth control. MEC was defined as the lowest concentration of lysin (± antibiotic) in a 2-fold dilution series that eradicated cells to below the limit of quantification (500 CFU/mL). Synergy was calculated using the fractional eradication concentration index; FECI = (MEC of agent A in combination/MEC of agent A alone) + (MEC of agent B in combination/MEC of agent B alone)^28^. The type of interaction was determined based on the median FEC index: synergy was defined as an FEC index ≤ 0.5, indifference as an FEC index > 0.5 to 4, and antagonism as an FEC index > 4.

## Simulated endocardial vegetation (SEV) assay

SEVs were generated following a modified version of Entenza et0al. (2009)^29^. For each reaction, 0.11mL of human plasma (pooled, sodium citrate anticoagulant; Dunn Labortechnik GmbH) was added to the wells of a reaction plate. The standardized inoculum corresponding to 10⁶ CFU per well was added, and input cell density was confirmed by serial dilution and quantification. Coagulation was triggered by adding 26 µL bovine thrombin (Sigma Aldrich; final concentration 10001U/mL), resulting in instantaneous clot formation. The resulting clots (mean mass 0.101±10.011g) were transferred into 5-fold excess volumes of cation-adjusted Mueller–Hinton broth containing 50% human plasma (caMHB–50% HuP) and incubated aerobically for 31h at 371°C with shaking at 650 rpm to promote bacterial proliferation within the fibrin matrix. Clots were washed 3x in 400 µL sterile DPBS by transfer and submersion using a pipette tip. After washing, clots were placed into deep-well plates containing 400 µL of antimicrobial solutions prepared in caMHB–50% HuP. Untreated clots served as controls. Clot–antimicrobial combinations were incubated anaerobically at 371°C for 241h at 6501rpm using sealed anaerobic incubation boxes with an anaerobic atmosphere generation pouch and a colorimetric indicator. After 24 h, clots were washed three times with 400 µL sterile DPBS. Clots were transferred into deep-well plates containing 400 µL tissue plasminogen activator (tPA; Boehringer Ingelheim Pharma GmbH & Co.KG; 128 µg/mL in DPBS) and incubated for 11h at 371°C with shaking at 650 rpm. Quantitative spotting of bacteria was performed by five 10-fold serial dilutions in DPBS and spotting 2 µL of the dilutions onto TSA plates (BD). Plates were incubated overnight at 37°C. CFUs were counted, and CFU/mL values were calculated based on dilution factors. Antimicrobial killing efficacy was reported as log₁₀ reduction relative to untreated control clots.

### In vitro phagocytosis assay

Overnight bacterial culture was stained with 10 nM BD Horizon™ CFSE (Carboxyfluorescein succinimidyl ester, 565082, BD Biosciences) in DPBS for 45 min and presented to THP-1 human monocytes in the presence of enzymatically inactive Fc(wt) or Fc(PGLALA) containing lysins or control antibody (anti-SpA^30^) with a MOI of 10. After 1 h incubation the cells were washed, fixed and the increase in basal phagocytosis was measured using FACS Lyric (BD Biosciences) as an increase in green fluorescent signal (mean fluorescence intensity of THP-1 cells).

### In vivo studies

Animal studies were performed at Evotec (UK) under UK Home Office Licenses and with local ethical committee clearance. All experiments were performed in dedicated BSL2 facilities. SPF mice (CD1 and C57BL/6J) used in this work were supplied by Charles River Laboratories (Margate, UK). After arrival, mice were acclimatised for 7 days and housed in IVC cages at 22 ± 1°C, 60 % relative humidity, maximum background noise of 56 dB and under 12-hour light/dark cycles. They had free access to food, sterile water, and were provided with aspen chip bedding. In some efficacy studies, animals were supplemented with wet food mush and HydroGel for additional hydration.

### In vivo PK studies

In PK studies, each test article was administered once as a slow bolus intravenous (IV) injection via a lateral tail vein in a dose volume of 5 mL/kg to a cohort of 3 mice. Following test article administration, approximately 20 µL of whole blood was collected from each mouse via the lateral tail vein in sampling tubes as per the dose/sampling schedule (mRNA: 1 h, 2 h, 4 h, 8 h, 12 h, 24 h, 48 h, 72 h, and 96 h; Recombinant protein: 5 min, 15 min, 30 min, 1 h, 2 h, 4 h, 10 h, 24 h, 48 h, 72 h, and 96 h). At the end of study (96 h), samples were collected via cardiac puncture under terminal isoflurane anesthesia. Plasma was separated by centrifugation at 10,000 x g for 5 min at 4°C. All samples were immediately frozen in liquid nitrogen and stored at -80°C.

### ELISA

Levels of Fc-LysM-CHAP and LysM-CHAP proteins in mouse plasma were quantified using a sandwich ELISA approach. In short, plates (Sigma Aldrich, CLS9018-100EA) were coated o/n at 4°C with 100 µL of anti-LysM-CHAP rabbit polyclonal antibody per well of (Genscript, U120LHF280-4) diluted to a concentration of 0.5 µg/mL in coating buffer (Sigma Aldrich, C3041-50CAP). Plates were washed 3x with 300 µL DPBS supplemented with 0.05% TWEEN-20 (DPBS-T) followed by a blocking step with 200 µL SuperBlock (Thermo Fisher Scientific, 37580) per well for 1 h at RT. Afterwards plates were washed again 3x as described above. Plasma samples were diluted 1:100 in SuperBlock, and further serially diluted (5-fold) in 1% mouse plasma (MoP) in SuperBlock. A standard dilution series in 1% MoP in Superblock was prepared in 2-fold dilutions from 256-0.25 ng/mL. 48 µL per well of sample or standard dilution was added to the plates and incubated for 1 h at RT, followed by three washes as described above. Bound analyte was detected using 100 µL of biotinylated anti-L0482 rabbit polyclonal antibody (Genscript, U120LHF280-4) diluted to 0.4 µg/mL in Superblock and incubated for 1 h at RT, followed again by three washes as described above. 100 µl per well of Poly-HRP streptavidin (Thermo Fisher Scientific, 21140) diluted 1:2500 in Superblock was added to the plates and incubated for 1 h at RT, followed by five washes as described above. After the last washing step, plates were incubated in washing buffer for 5 min at RT. For detection, 100 µL per well of TMB solution (Sigma Aldrich, T0440) was added and incubated for 2-5 min followed by addition of 100 µL per well of 2 N sulphuric acid to stop the reaction. Signal was recorded at 450 nm in a Sunrise plate reader (Tecan). Signal from the standard curve was fit in GraphPad PRISM to a sigmoidal curve (4PL) and used to interpolate the lysin concentration in the plasma samples.

Plasma concentration data was analysed using GraphPad (version 10.2.3). Mean concentrations were used for calculation of the area under the concentration time curve (AUC).

### mRNA synthesis and formulation

DNA sequences were codon-optimized prior to gene synthesis. Transcribed RNA strand contains common structural elements optimized for stability and translational efficiency (CC413 as 5’-cap, AGA-dEarI-hAg as 5’-untranslated region [UTR], FI element as 3’-UTR, and a poly[A] tail measuring 100 adenosines with a linker at position 30 [A30L70])^31^ ^32^.

In vitro transcription of the RiboLysin performed as previously described^33^. The resulting 5’, N1-methylpseudouridine (m1Ψ)-capped RNA was isolated with magnetic beads, purified with cellulose, diluted in RNAse-free water and stored in nuclease-free reaction tubes (Eppendorf). The transcribed RNA was quality-assessed (Agilent 2100 Bioanalyzer, Agilent Technologies Inc.) and quantitated (Nanodrop 2000c, Thermo Fisher Scientific) prior to encapsulation in LNP (ionizable cationic lipid, a polyethylene glycol (PEG) lipid, phospholipid, and cholesterol) followed by storage at -80°C. The formulations were tested for particle size, distribution, encapsulation efficiency, and RNA concentration. A similar LNP formulation was tested previously in vitro and in vivo^34^.

### In vivo efficacy

Inoculum preparation and quantitative burden in mouse tissue. Infections were performed with freshly prepared Sa (ATCC43300, NCTC 8178 or BAA-1717) grown to logarithmic phase. The target inoculum was confirmed by quantitative plating on mannitol salt agar (MSA) plates. Mice were euthanized by terminal cardiac bleed under isoflurane anesthesia and death confirmed by cervical dislocation. Tissue samples were homogenized in DPBS using Precellys homogenizer and homogenates were serially diluted 1:10 in sterile DPBS, followed by quantitative plating on MSA plates. Plates were incubated at 37°C for 18-24 h and bacterial colonies enumerated.

Neutropenic mouse thigh infection model. Neutropenia was induced in male C57BL/6J mice (n=5) with cyclophosphamide (subcutaneous (SC), 150 mg/kg 4 days before infection and 100 mg/kg 1 day before infection) and animals were infected with intramuscular injection (IM) of 5x10^3^ CFU/thigh Sa (ATCC43300) in both thighs. Recombinant lysins were applied 1 h post infection (hpi), intravenously (IV), every 12 h (q12), at dose of 500 µg/mouse. Vancomycin was used as a comparator drug at dose of 100 mg/kg, q12, treatment initiated 1 hpi.

### Quantitative bacterial burden in each thigh was determined at 25 hpi (pre-treatment at 1 hpi)

Mouse IV sepsis model. Mice were infected with a 100 µL injection in the lateral tail vein of 1x10^7^ or 5x10^7^ Sa (NCTC 8178 or BAA-1717). Body weights were recorded and weight loss relative to day 0 calculated over the course of the study (ethical limit >20% weight loss). The clinical condition of mice was observed and scored regularly throughout the study to monitor animal welfare. In mRNA/LNP studies, test articles were administered IV once at 1 hpi and at doses of 2, 0.5 or 0.2 mg/kg. Vancomycin treatment (25 mg/kg, IV, q12 h from 1 hpi) was used as comparator. In recombinant protein studies, lysins were administered at dose of either 17.5 mg/kg or 3.5 mg/kg, starting from 1 or 13 hpi, q12 or q24. Vancomycin was used as a comparator drug at dose of 25 mg/kg, q12, from 1 or 13 hpi, alone or in combination with tested lysins. Quantitative bacterial burden in both kidneys together was determined at 72 hpi (pre-treatment at 1 hpi).

---

**Extended Data Figure 1.**
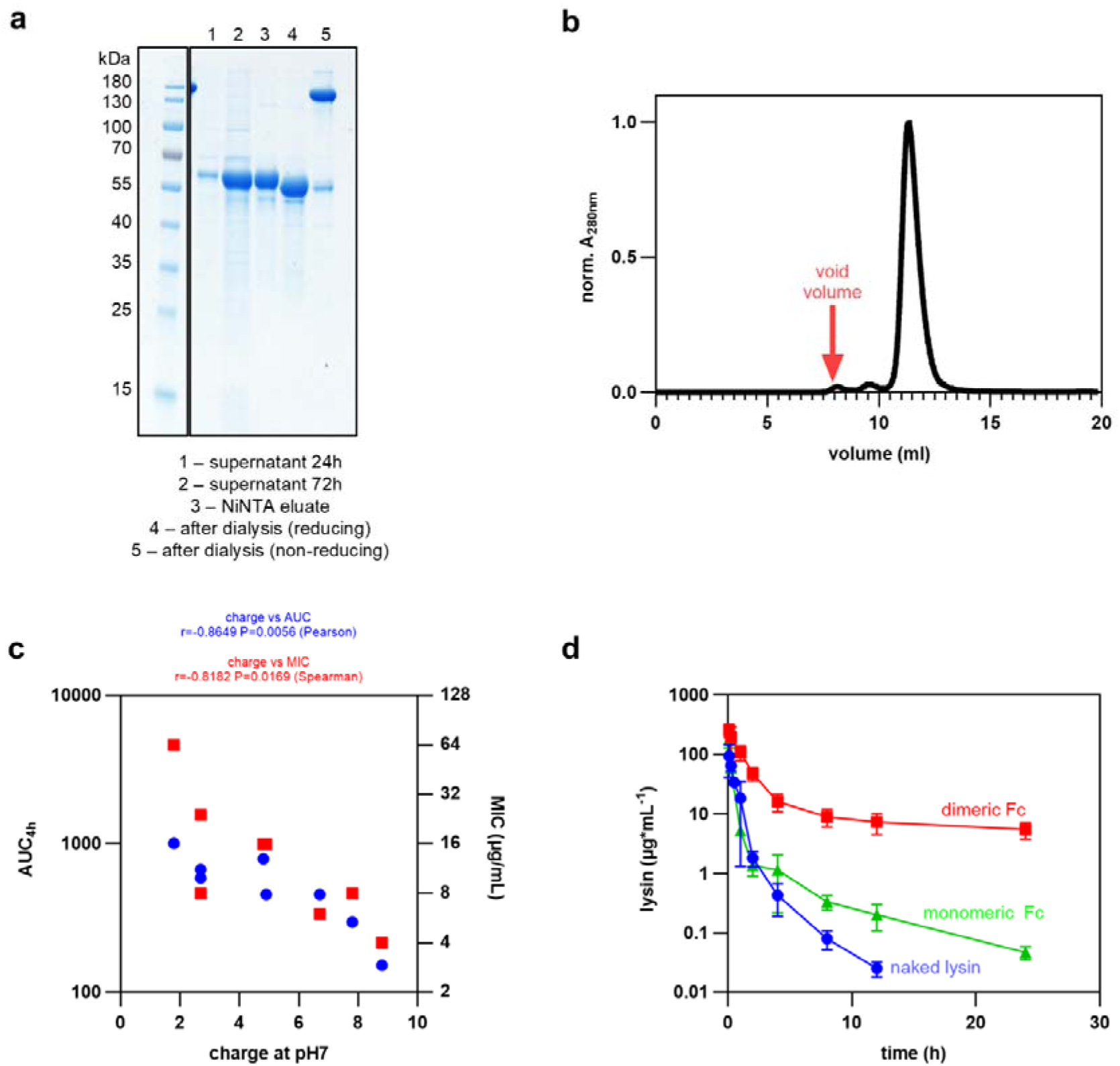
: a.) Purification of Fc-hel8-L1-3. Fc-hel8-L1-3 was purified from Expi293 supernatants by Ni-NTA affinity chromatography, followed by removal of the purification tag, protein concentration and buffer exchange. Samples of culture supernatants at 24 h and 72 h post transfection, Ni-NTA eluates and samples after dialysis were taken and analyzed by SDS-PAGE under reducing and (where indicated) non-reducing conditions followed by Coomassie staining. b.) SEC behavior of Fc-hel8-L1-3. Dialyzed samples of purified Fc-hel8-L1-3 were analyzed by size exclusion chromatography (SEC) using a Superdex 200 increase 10/300 gl column. Protein absorbance at 280nm was normalized against the maximum signal recorded during the SEC run. The void volume of the column at ∼8 mL is indicated. c.) Impact of protein charge on PK and MIC of different Fc-LysM-CHAP variants. AUC at 4 h (AUC_4h_, blue) was determined from the PK curves shown in Figure 1c. MIC values (red) are as determined in Figure 1b. AUC_4h_ (log10 scale) and MIC (log2 scale) were plotted against the charge at pH7 of each Fc-LysM-CHAP variant (as determined from the amino acid sequence). The relationship between charge and AUC was tested for linear correlation with the Pearson method; the relationship between charge and MIC was tested for non-linear correlation with the Spearman method (correlation coefficients (r) and significance values (P) are indicated). AUC: area under curve; MIC: minimal inhibiting concentration. d.) PK comparison of dimeric and monomeric Fc fusions. PK curves of dimeric (red) and monomeric (green) formats of Fc-hel8-L1 and the corresponding naked lysin (L1, blue) were determined as described in Figure 1c.). Datapoints represent mean and standard deviation (n=3).

**Extended Data Figure 2.**
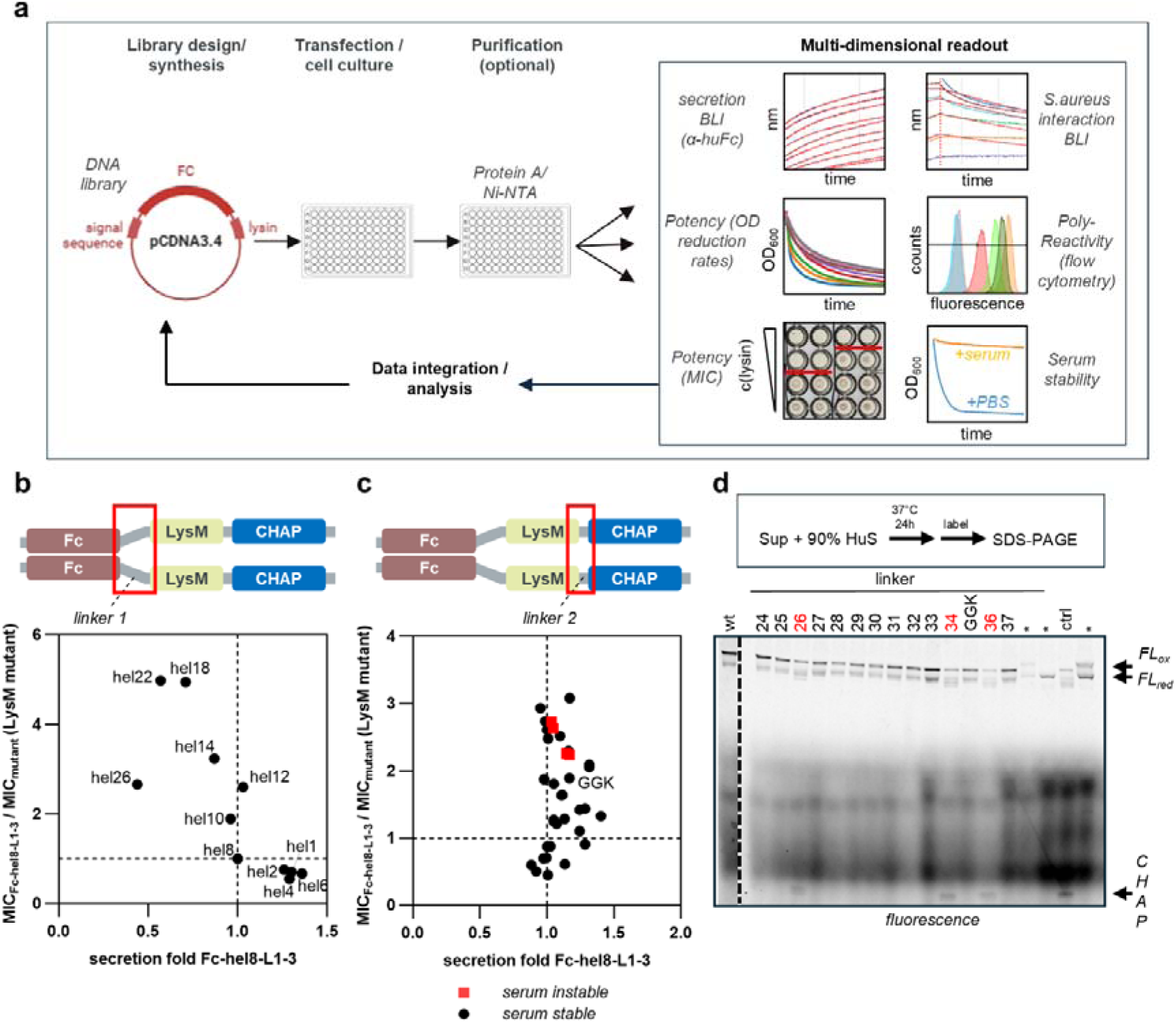
: a.) Summary of the Lysin screening platform. A DNA-library encoding lysin derivatives is transfected into Expi293 cells. Supernatants containing secreted lysins are harvested, optionally purified and lysin concentrations determined by biolayer interferometry. Lysins are further subjected to biochemical characterization, including assays for measurement of ODRRs, MIC, Sa binding, poly-reactivity and serum stability (see Supplemental Discussion). Data is integrated and used to design the next library for screening. α-huFc: BLI-sensor tips decorated with an antibody specific for the human Fc domain. pCDNA3.4: DNA vector for library assembly. b.) Effect of helical linker length on lysin MIC. As described in Figure 2c but with lysin activity quantified with MIC. Supernatants were serially diluted and mixed with 5x10 CFU/mL Sa BAA-1717, followed by incubation at 37 °C for 18 h. Bacterial growth was assessed by optical density at 600nm (OD). MIC was determined as the lowest lysin concentration at which OD was below the threshold determined for a sterile control condition. MICs were normalized relative to the Fc-hel8-L1-3 construct (indicated by vertical and horizontal dotted lines) using the formula MIC_Fc-hel8-L1-3_/MIC_variant_. Secretion of each variant in supernatants was measured by BLI and normalized to the secretion of the parental Fc-hel8-L1-3 (“secretion fold Fc-hel8-L1-3”). c.) Effect of linker-2 amino acid sequence on lysin MIC. As in b.) but with a set of Fc-hel8-L1-3 derivatives containing different sequences in the linker between the LysM and CHAP domains. Supernatants were further analyzed for stability in the presence of human serum (see panel d). Variants which were not stable in serum for >24 h at 37 °C are highlighted in red. Secretion and activity of parental Fc-hel8-L1-3 are indicated by vertical and horizontal dotted lines, respectively. Secretion and activities of the shortlisted GGK linker is indicated (“GGK”). d.) Serum stability of Fc-hel8-L1-3 variants containing different linker-2 sequences. Supernatants from the experiment described in panel c were mixed 1:9 with human serum and incubated anaerobically at 37 °C for 24 h. Lysins were labeled with 1 µM Pep1a-CMK for 5 min, followed by addition of non-reducing sample buffer and SDS-PAGE. Lysin fragmentation was then assessed by fluorimetry. Full-length Fc-LysM-CHAP migrates as an oxidized (FL_ox_) and reduced (FL_red_) species under non-reducing conditions. Migration of the free CHAP domain released by serum proteases is indicated. Serum u nstable constructs are highlighted in red. ctrl: Fc-LysM-CHAP constructs containing the serum-sensitive linker-1 sequence of LytN. wt: Fc-hel8-L1-3. The lead construct containing the GGK linker is indicated.

**Extended Data Figure 3.**
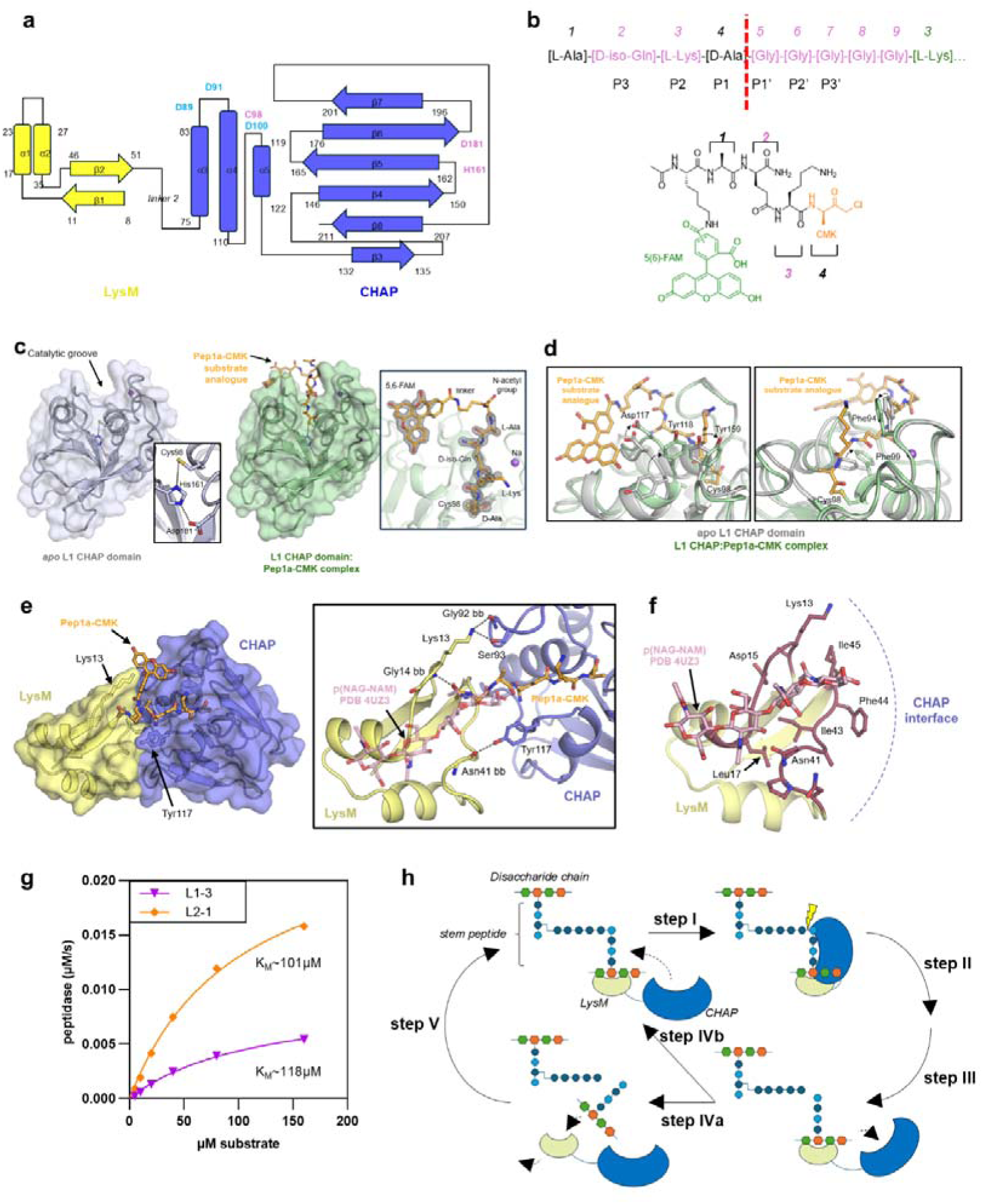
: a.) Secondary structure of LysM-CHAP derivatives. α-helices (α1- α5) are indicated as boxes, β-sheets (β1- β7) as arrows. LysM (yellow) and CHAP (blue) domains are indicated. Numbers indicate boundaries of secondary structure elements. Residues involved in the catalytic triad are highlighted in pink and labeled in single letter amino acid code. Residues involved in coordination of the Ca /Na ion are highlighted in turquoise. b.) Structure of the Sa stem peptide and its analogue Pep1a-CMK. (top) Amino acid sequence of the Sa stem peptide. Numbers indicate positions of amino acids as referred to in the Supplemental Discussion. Amino acids highlighted in pink are variable between bacterial species. Protease subsites are indicated below the stem peptide sequence according to Schechter & Berger^1^. (bottom) Structure of Pep1a-CMK. Stem peptide amino acids are shown in black. Correspondence to positions in the top panel are indicated. The C-terminal alanyl-chloromethyl ketone (CMK) is shown in orange. The fluorescein fluorophore attached via an N-terminal lysin residue is shown in green. c.) Comparison of apo- and Pep1a-CMK bound holo L1 structures (PDBs: 11CF and 11CH, respectively). (left) The L1 CHAP domain in the apo form is shown as a cartoon (light blue). The inset shows ball-and-stick models of the residues which coordinate the catalytic cysteine at position 98. (right) The L1 CHAP domain bound to Pep1a-CMK is shown as a green cartoon. Pep1a-CMK is shown as a ball-and-stick model in orange. The inset shows the electron density map of Pep1a-CMK as a mesh. Stem peptide amino acids and the 5,6-FAM moiety are indicated. d.) Side chain rearrangements upon substrate binding in the L1 CHAP domain. Substrate-free apo- (grey; PDB: 11CF) and Pep1a-CMK-bound holo- (green; PDB:11CH) L1 structures were structurally aligned. Left: Zoom-in on the conformations of side chains of Asp117, Tyr118 and Tyr159. Right: Zoom-in on side chain conformations of Phe94 and Phe99. e.) Molecular interactions between substrate, LysM and CHAP domains. Left: Surface and cartoon representation of the interaction between LysM (yellow) and CHAP (blue). Residues Lys13 and Tyr117 are indicated as ball-and-stick models. Right: Zoom in on residues involved in contacts between LysM (yellow) and CHAP (blue) domains. Interacting residues are shown as stick models. Red dashed lines indicate interacting amino acids pairs. Pep1a-CMK is shown as an orange ball-and-stick model. The NAG-NAM chain of PDB: 4UZ3 is shown as a pink ball-and-stick model. bb: interactions involving the amino acid backbone. p(NAG-NAM): NAG-NAM polymer. f.) LysM residues in proposed contact with the NAG-NAM polymer. Shown is a magnified view of the LysM-domain with a modelled carbohydrate chain (shown as pink ball-and-stick model). Residues in proximity to the modelled carbohydrate chain (pink) are shown as sticks and colored in brown. The interface to the CHAP domain is indicated with a dotted arc. g.) Peptidase activity of L1-3 and L2-1 at different substrate concentrations. Peptidase activity was quantified as the maximum rate of cleavage of a fluorogenic stem peptide substrate analogue (Pep1a) at different concentrations and at a fixed lysin concentration of 2.5 µM. Datapoints were fit to the Michaelis-Menten model to estimate the Michaelis-Menten constant K_M_ values for the two variants. h.) Proposed mode of action of LysM-CHAP. Arrows indicate reaction steps, dotted arrows indicate domain movements. LysM-CHAP binds to a peptidoglycan cross-bridge via interaction between the carbohydrate chain and the LysM-domain. LysM binding increases the k_on_ for CHAP binding to the stem peptide (step I). Once bound, the CHAP domain cleaves the stem peptide (step II). After cleavage, the CHAP domain has a reduced affinity to the hydrolyzed stem peptide (i.e. the reaction product), promoting product dissociation (step III). Dissociation of the CHAP domain decreases the affinity of LysM binding to the carbohydrate chain, causing full dissociation of LysM-CHAP from the cleaved cross-bridge. The unbound LysM domain now allows LysM-CHAP to engage with the next cross-bridge (step IVa). Alternatively, the LysM domain slides on the carbohydrate chain to the next cross-bridge (step IVb).

**Extended Data Figure 4.**
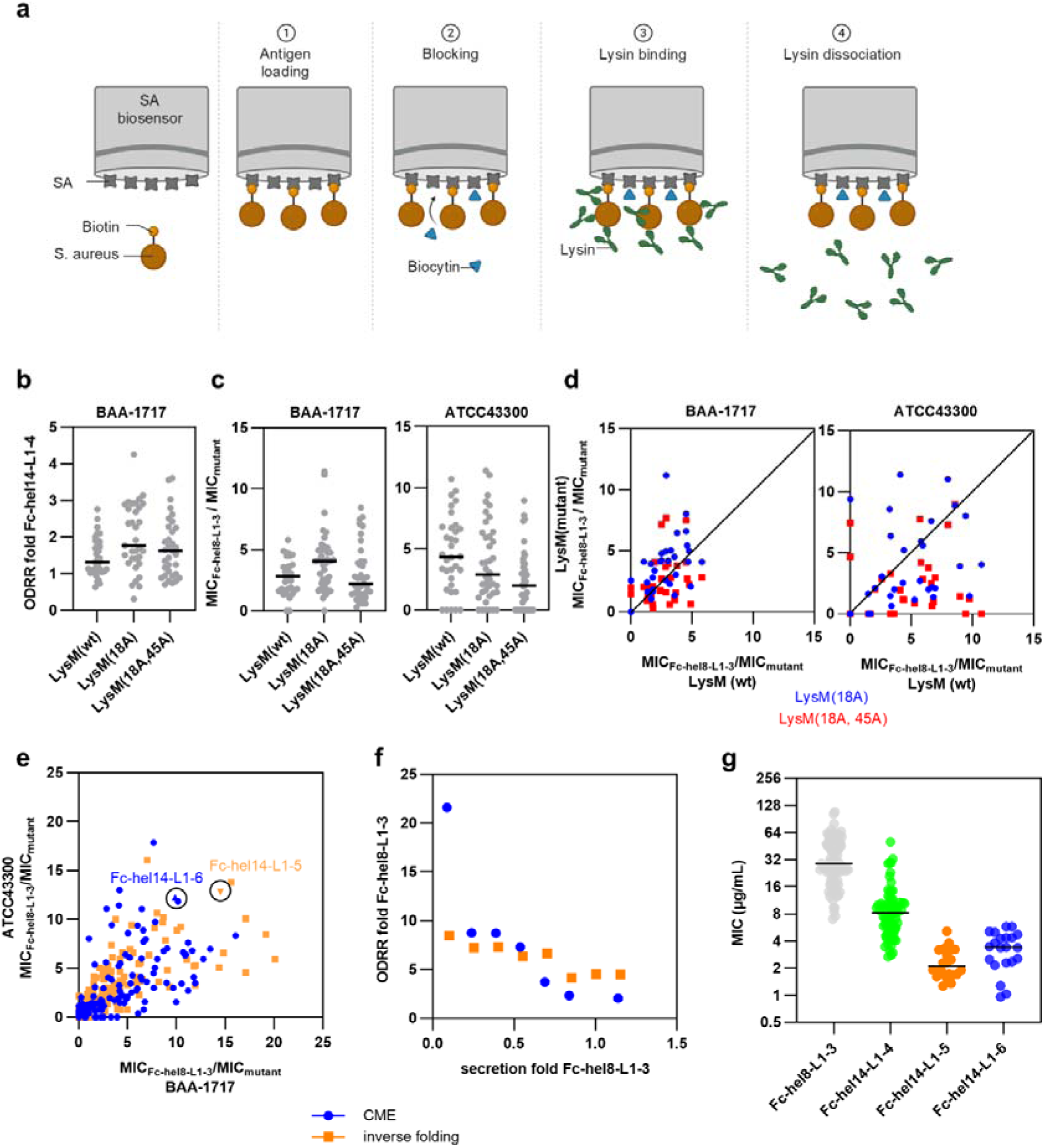
: a.) Principle of the BLI assay for measuring binding between lysins and surface immobilized Sa cells. Biotinylated Sa cells are immobilized on streptavidin (SA) biosensors (step 1), followed by blocking with biocytin (step 2). Biosensors are then exposed to solutions containing different Fc-LysM-CHAP variants at increasing concentrations, followed by measurement of association rates (k_on_; step 3). After binding, the sensors are transferred to buffer solutions, followed by measurement of lysin dissociation rates (k_off_; step 4). b.) Impact of LysM mutations on ODRRs of active site variants of Fc-hel14-L1-4. ODRRs of a set of 33 combinatorial motif engraftment (CME) active site derivatives of Fc-hel14-L1-4 containing either the wild type LysM domain (LysM(wt)), or LysM domains harboring the S18A (LysM(18A)) or S18A/I45A (LysM(18A,45A)) mutations were determined in supernatants and normalized to parental Fc-hel14-L1-4. Shown are normalized ODRRs of individual CME variants. Horizontal lines indicate means of all CME variants in the respective LysM background. c.) Impact of LysM mutations on MICs of active site mutants of Fc-hel14-L1-4. Supernatants described under b.) were subjected to growth inhibition assays against Sa strains BAA-1717 (left) and ATCC 43300 (right). MICs were normalized relative to the Fc-hel8-L1-3 construct using the formula MIC_Fc-hel8-L1-3_/MIC . Shown are values for individual variant constructs and means of all CME variants in each LysM background. d.) Effect of LysM mutations on MICs of active site mutants of Fc-hel14-L1-4. MIC data from panel c.), but with fold Fc-hel8-L1-3 MIC changes of LysM(wt) variants plotted against the corresponding LysM(18A) (blue) and LysM(18A,45A) (red). MIC data against Sa BAA-1717 (left) and ATCC 43300 (right) are shown. e.) High throughput screen for active site mutants of Fc-hel14-L1-4(18A) with improved MICs. Supernatants described in Figure 3e were subjected to growth inhibition assays against Sa BAA-1717 (X-axis) and ATCC 43300 (Y-axis). Determined MICs were normalized relative to the Fc-hel8-L1-3 construct using the formula MIC_Fc-hel8-L1-3_/MIC. CME variants are shown in blue, inverse folding variants are shown in orange. Lead variants Fc-hel14-L1-5 and Fc-hel14-L1-6 are highlighted. f.) Trade-off between secretion and ODRR for variants designed by CME and inverse folding approaches. Same as Figure 3e.), but only showing secretion and activities of the most active CME (blue) and inverse folding (orange) variants at each secretion bin. g.) Summary of MICs of Fc-hel8-L1-3, Fc-hel14-L1-4, Fc-hel14-L1-5 and Fc-hel14-L1-6 as determined from supernatants. Shown are individual MIC values determined for Sa BAA-1717 from independent transfections. Median MICs are indicated with horizontal lines.

**Extended Data Figure 5.**
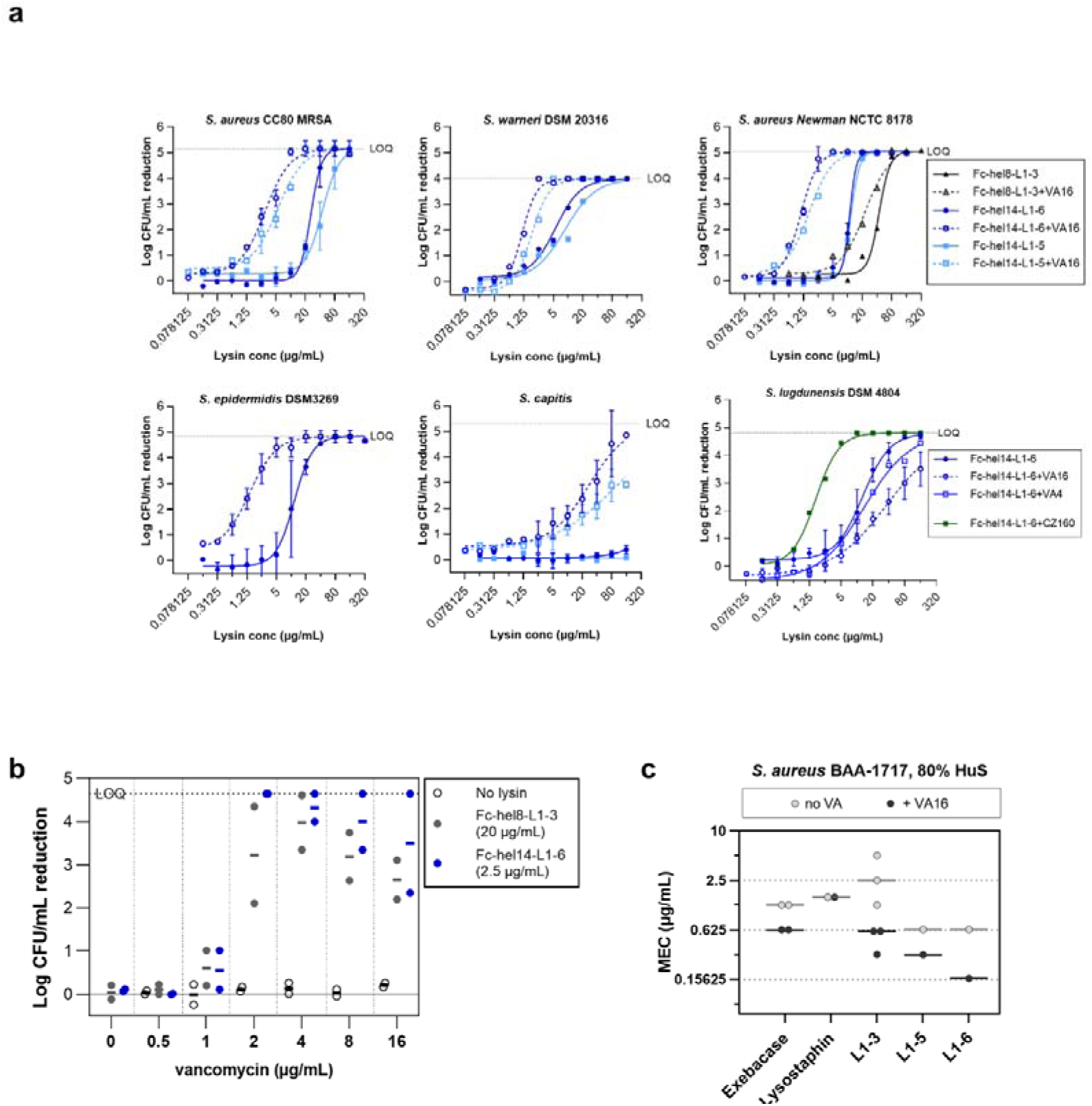
: a.) Activity of lysins with or without antibiotics in 80% human serum (HuS). Lysin dose responses were tested alone or with 16 or 4 µg/mL vancomycin or 160 µg/ml cefazolin on 5×10 to 2×10 CFU/mL of different staphylococcal strains for 3 h in 80% HuS. Mean and standard deviation from independent experiments is shown. Mean maximum log reduction (LOQ) is indicated as dashed horizontal line. 4PL curves were fitted to the data with the top constraint to mean LOQ. Treatment with antibiotic alone was within ±0.5 log_10_ CFU/mL reduction of all strains. LOQ: limit of quantification. b.) Vancomycin titration in combination with lysis. Fc-hel8-L1-3 and Fc-hel14-L1-6 were tested at constant concentrations in combination with 0-16 µg/mL vancomycin on Sa BAA-1717 in 80% HuS. Log CFU/mL reduction was quantified after 3 h relative to the untreated control. Data from two independent experiments are shown, mean is indicated as a horizontal line. c.) MEC with and without vancomycin of non Fc-tagged lysins of different architectures. Lysin dose responses were tested on 5×10 to 2×10 CFU/mL Sa cells as single (grey circles) or combination treatment with vancomycin at a fixed concentration of 16 µg/mL (black circles).

**Extended Data Table 1.**
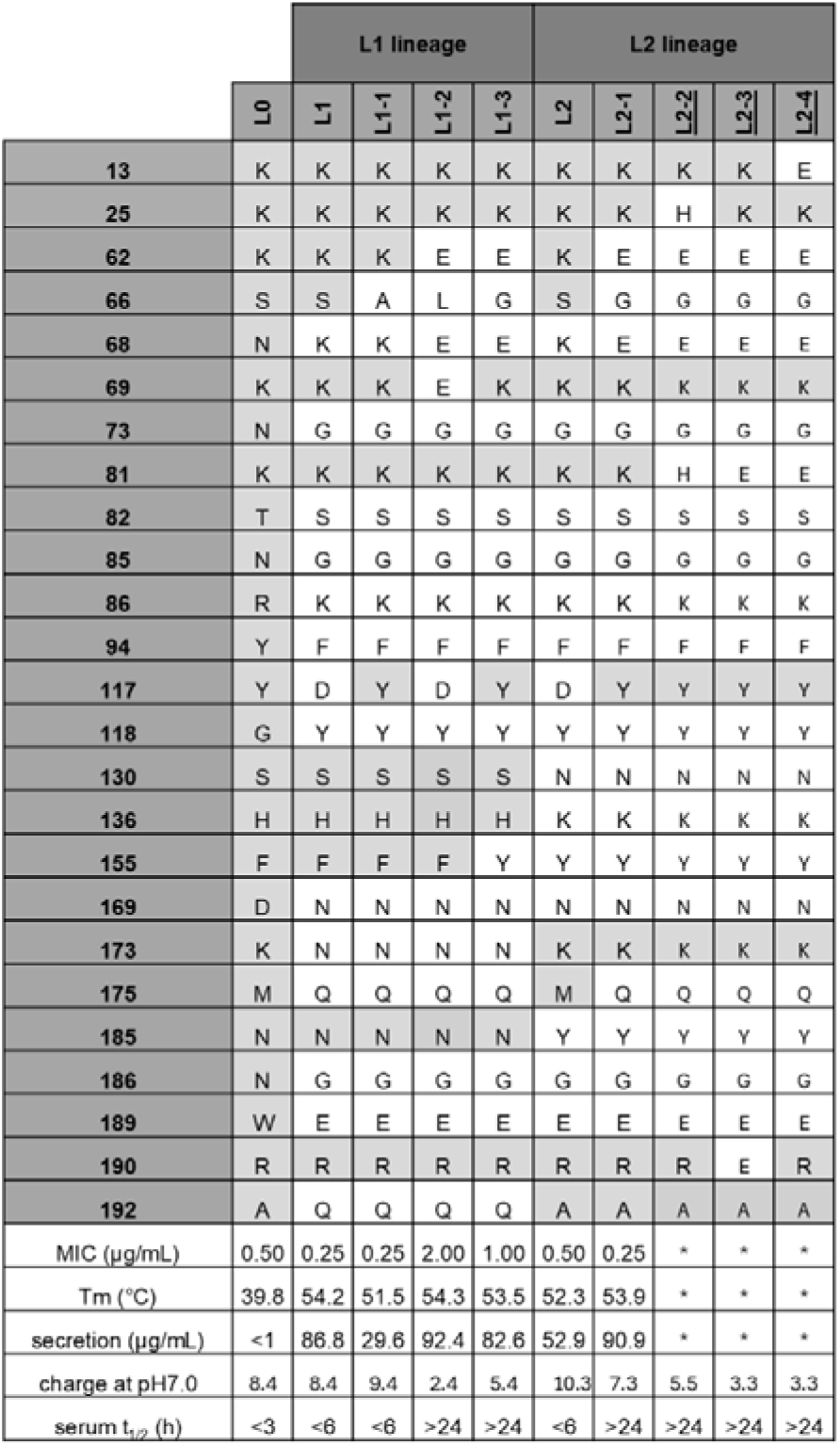
: Comparison of engineered LysM-CHAP variants. Numbers indicate amino acid positions at which variants differ. Amino acids found at these positions in the individual variants are shown in single letter format. Light grey boxes indicate amino acids found in the wild type LytN sequence. MIC against Sa ATCC 43300, melting temperature (Tm) as measured by NanoDSF, secretion and half-life at 37°C in 90% human serum (serum t_1/2_). Underlined lysins were only tested as Fc-fusion proteins. Asterisks (*) indicate that biochemical data was not recorded for naked LysM-CHAP versions (only for Fc-fusions).

**Extended Data Table 2.**
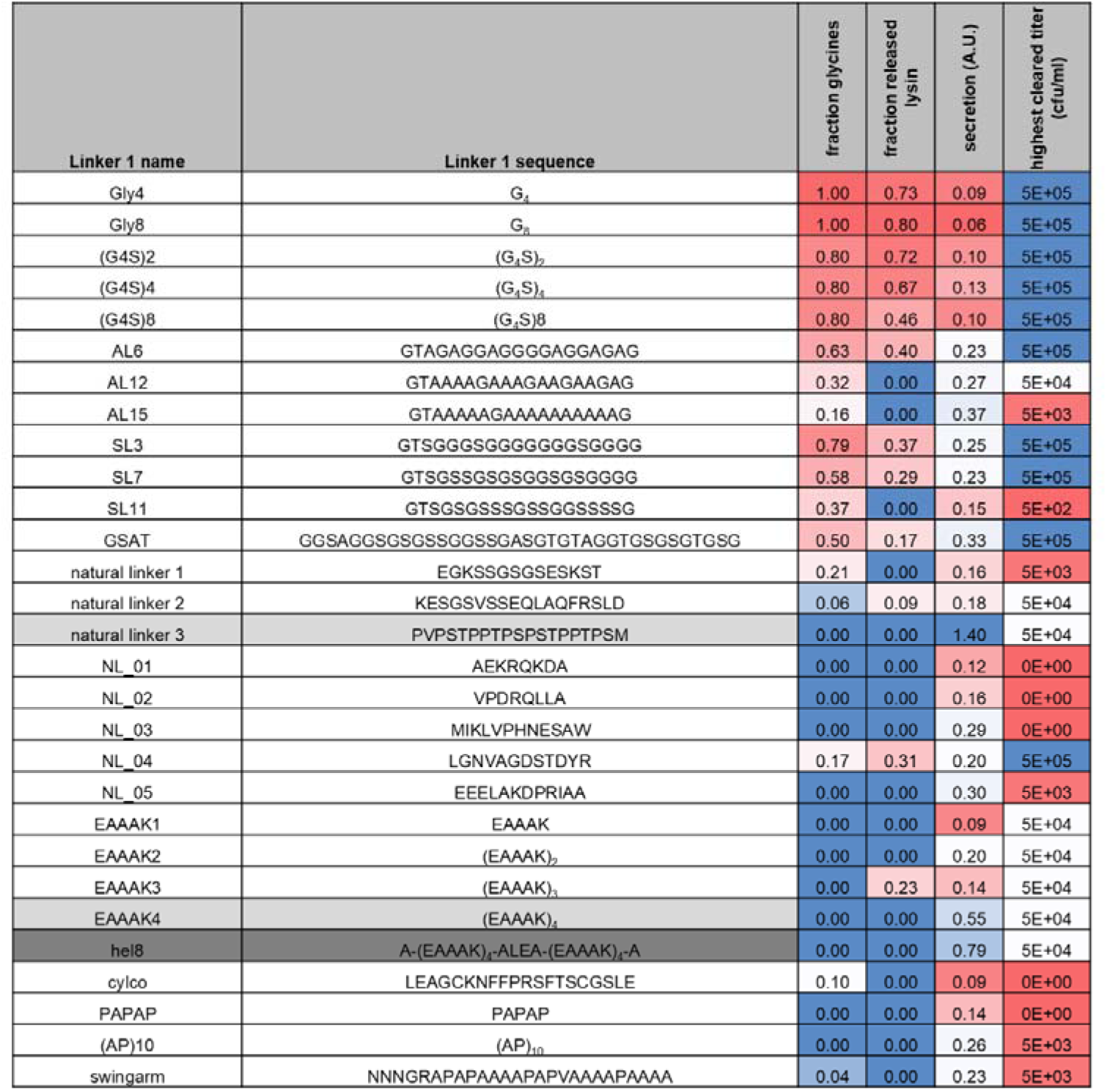
Screen for linker-1 in Fc-LysM-CHAP. Human IgG1-Fc domain was fused to the N-terminus of LysM-CHAP variant L1 via different linkers (linker-1 name and amino acid sequence in single letter code shown in the first and second column, respectively). Constructs were transfected into Expi293 cells and supernatants containing secreted protein tested for protein content and integrity by SDS-PAGE followed by Coomassie staining and for activity using growth inhibition assays with different colony forming units (CFU) input of Sa ATCC 43300. The fraction of glycine residues (column 3) was determined from the amino acid sequence of the linkers. Bands on SDS-PAGE were quantified to determine the fraction of free lysin (column 4) and the total amount of full length secreted lysin. The secretion level (column 5) was determined by normalizing the total amount of full length lysin by a L1 control loaded on the same gel. Activity was quantified by determining the highest CFU input of Sa that was cleared by the respective supernatant. Linkers with viable secretion and activities are highlighted in grey. The hel8-linker is highlighted in dark grey.

**Extended Data Table 3.**
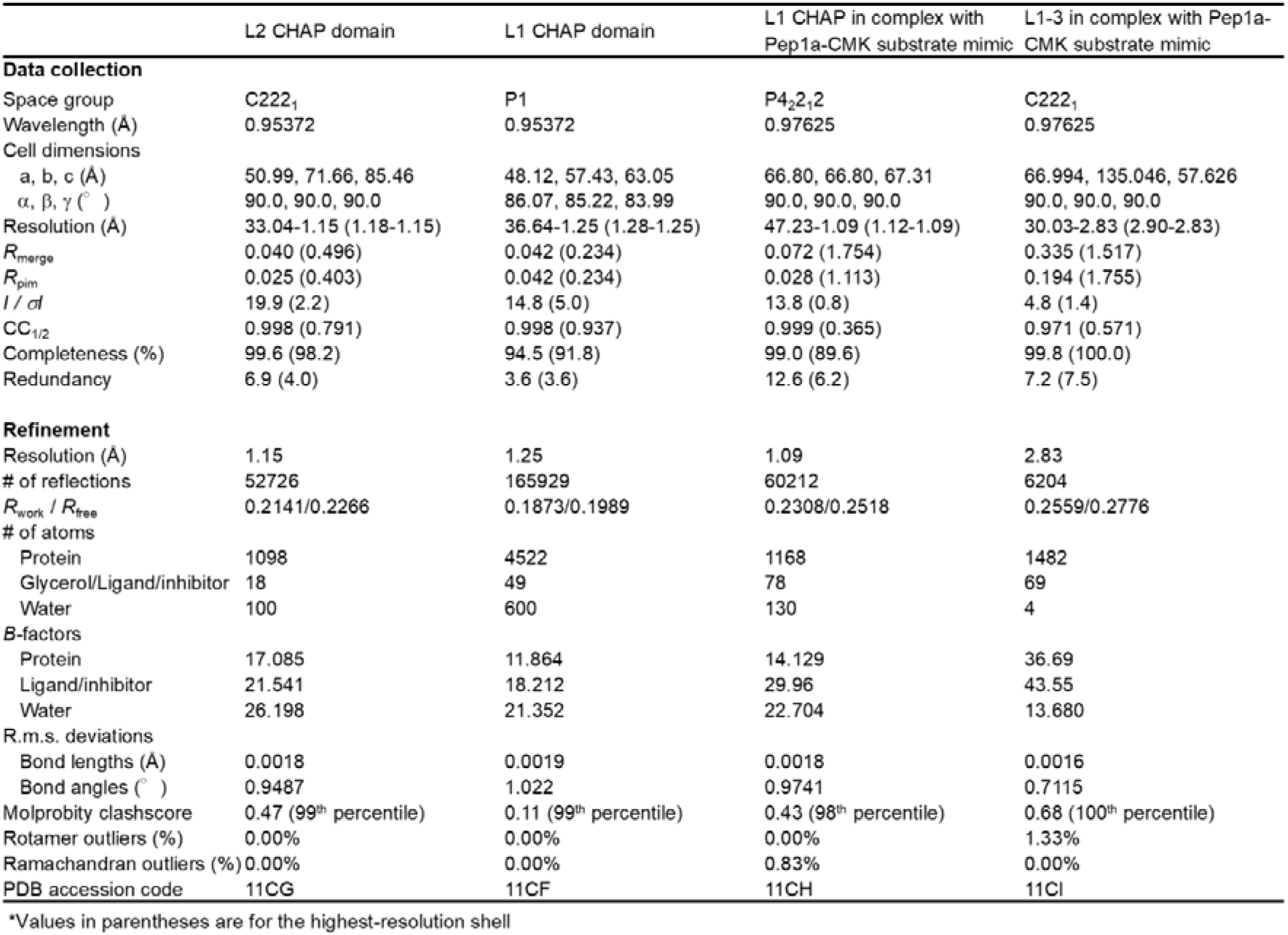
X-ray crystallographic data collection and refinement statistics. *Values in parentheses are for the highest-resolution shell.

**Extended Data Table 4.**
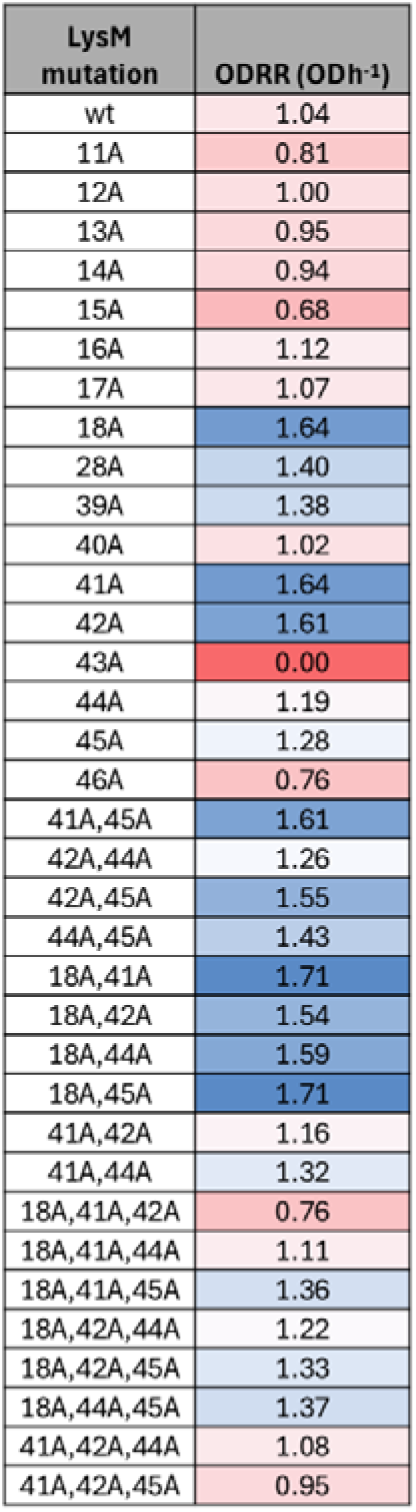
: ODRRs of LysM mutants of Fc-hel14-L1-4 and combinations thereof. Variants of Fc-hel14-L1-4 containing single mutations in the LysM domain and indicated combinations thereof were purified and subjected to OD reduction assays against Sa ATCC 43300 at a final enzyme concentration of 0.2 µM. ODRRs for each variant are color-coded blue - high activity, red – low activity).

| LysM mutation | ODRR (ODh<sup>-1</sup>) |
| --- | --- |
| wt | 1.04 |
| 11A | 0.81 |
| 12A | 1.00 |
| 13A | 0.95 |
| 14A | 0.94 |
| 15A | 0.68 |
| 16A | 1.12 |
| 17A | 1.07 |
| 18A | 1.64 |
| 28A | 1.40 |
| 39A | 1.38 |
| 40A | 1.02 |
| 41A | 1.64 |
| 42A | 1.61 |
| 43A | 0.00 |
| 44A | 1.19 |
| 45A | 1.28 |
| 46A | 0.76 |
| 41A,45A | 1.61 |
| 42A,44A | 1.26 |
| 42A,45A | 1.55 |
| 44A,45A | 1.43 |
| 18A,41A | 1.71 |
| 18A,42A | 1.54 |
| 18A,44A | 1.59 |
| 18A,45A | 1.71 |
| 41A,42A | 1.16 |
| 41A,44A | 1.32 |
| 18A,41A,42A | 0.76 |
| 18A,41A,44A | 1.11 |
| 18A,41A,45A | 1.36 |
| 18A,42A,44A | 1.22 |
| 18A,42A,45A | 1.33 |
| 18A,44A,45A | 1.37 |
| 41A,42A,44A | 1.08 |
| 41A,42A,45A | 0.95 |

