## Supplemental Discussion for "Multi-dimensional optimization of a lysin towards a ribolysin against life-threatening *S. aureus* infections: Fc-LysM-CHAP and its strong synergy with standard of care antibiotics"

**Lysin screening platform**

The Lysin screening platform is summarized in Extended Data Fig. 2a and is centered on a pCDNA3.4-derived backbone vector which drives gene expression under the constitutive CMV promoter. The plasmid encodes the fusion of a mammalian signal sequence (a mouse IgKappa signal sequence mutant), a cleavable purification tag, and the human dimeric IgG1-Fc domain, followed by a multiple cloning site into which LysM-CHAP variants can be cloned rapidly and in high throughput. DNA libraries containing different Fc-LysM-CHAP variants are normalized for concentration, arrayed with control constructs into 96 well plates and then mixed with transfection reagents using a robotic pipetting platform. Constructs are transfected into Expi293 suspension cell cultures grown in multi-well plates and supernatants harvested for functional testing after 72 h of culture.

Our platform allows assessment of lysin properties directly in supernatants or after purification. For the latter, we adapted 24-well and 96-well formats of nickel-nitrilotriacetic acid (Ni-NTA)- or protein A affinity matrices, which can be used to isolate, wash and elute secreted lysins from supernatants via their N-terminal hexa-histidine tags or via their Fc-domain, respectively.

The presence of a human Fc domain in all constructs allows us to further determine the secreted lysin titer in high throughput and accuracy using biolayer interferometry (BLI) combined with biosensor tips that recognize the human Fc domain.

For measurement of lysin potency, optical density (OD) reduction assays are performed in which the rate of OD reduction at 600 nm of a bacterial *Sa* solution is quantified from the maximum slope of the OD reduction curve. The OD reduction rate (ODRR) of a lysin variant at the determined concentration can be compared relative to the ODRR of a control lysin at the same concentration to determine a fold change in activity, which can be used to rank lysins for potency.

As a complementary activity measurement, we use an automated liquid handling platform to perform a total of 8 two-fold serial dilutions of the lysins followed by mixing with a *Sa* solution containing a defined concentration of bacterial cells. The dilution plates are incubated for ~18 h at 37°C and the OD of each sample and dilution is measured in a plate reader. This data allows us to identify the minimum dilution of a given lysin required to prevent the outgrowth of the bacterial suspension. In combination with the lysin concentration assessed by BLI the minimal inhibitory concentration (MIC) of the lysin can be determined.

Depending on the screening purpose, the lysin samples can be subjected to additional biochemical assays, which we have similarly adapted to the format of our screening platform. These assays allow us to quantify the kinetics of lysin binding to different *Sa* strains immobilized on biosensors by BLI, the assessment of poly-reactivity of a lysin variant to mammalian cell surfaces by flow cytometry and the assessment of serum stability by SDS-PAGE or OD reduction assays after incubation in different biological matrices.

We have automated data extraction, quality control, quantification and normalization of biochemical parameters from each of these assays using Python scripts, which allow us to assign multidimensional properties to each amino acid sequence in a standardized form with minimal user input. These datasets are then used to extract design principles for e.g. lysin potency and guide the generation of lysin amino acid sequences for the next screening library.

**Structural analysis of LysM-CHAP derivatives using X-ray crystallography**

We were initially unable to crystallize full-length versions of LysM-CHAP variants, likely because the flexible linker between LysM and CHAP allows the two domains to exist in variable conformations. We therefore focused on structural analysis of isolated CHAP domains (amino acids 74-215). Crystallization screens in the absence of ligands yielded crystals for the L1 and L2 CHAP domains which were used to determine structures at 1.18 Å and 1.28 Å resolution, respectively (Extended Data Table 3). L1 and L2 display the typical globular CHAP-architecture seen in the previously determined X-ray structures of LysK^1,2^ and LysGH15^3^, with a two-lobe fold formed by three N-terminal alpha-helices packed against six C-terminal beta-sheets (Extended Data Fig. 3a). The two lobes form a pocket which contains the catalytic triad formed by Cys98, His161 and Asp181. The catalytic cysteine is positioned at the N-terminus of the second alpha-helix in the CHAP domain (α4 in Extended Data Fig. 3a) near a metal-binding motif formed by residues Asp89, Asp91 and Asp100. Due to the crystallization in presence of high concentrations of NaCl, the motif binds Na^+^ in our structures but likely coordinates Ca^2+^ under physiological conditions (Fig. 3b).

Pilot studies, in which different CHAP domains had been crystallized in presence of a full-length synthetic *Sa* stem peptide (L-Ala-isoD-Gln-L-Lys-L-Ala-(Gly)_5_) had failed to resolve the substrate in the catalytic pocket of the CHAP domain, likely due to cleavage during crystallization or low too low affinity (data not shown). To stabilize the CHAP-substrate complex, we designed a modified stem peptide (Pep1a-CMK, Extended Data Fig. 3b) in which the amino acid sequence starting at the P1’ site (GGGGG) was substituted with a chloro-methyl ketone (CMK) group, which would form a covalent bond to the catalytic Cys98 upon substrate engagement and trap the complex in a conformation close to the transition state during peptide hydrolysis. The substrate was further appended with a FAM moiety for fluorescent detection.

Purified L1 CHAP domain was labelled with an excess of Pep1a-CMK followed by removal of unreacted substrate by gel filtration and crystallization. This approach allowed us to gather crystals of substrate-bound L1 CHAP and solve its structure at 1.08 Å resolution. The high overall resolution of the structure and the fact that substrate atoms were clearly resolved allowed us to model the interactions between the substrate and the catalytic pocket in atomic detail (Fig. 3b and Extended Data Fig. 3c).

The structure shows Pep1a-CMK bound between the two lobes of the catalytic pocket, with Cys98 forming a covalent adduct to the remaining carbon of the CMK moiety. In the stem peptide-bound conformation, the sulfur atom of Cys98 is 2.6 Å away from the carbon of the D-Ala carboxyl group (Position 4 in Extended Data Fig. 3b). This distance would need to decrease by 0.8 Å during hydrolysis of a native substrate, a conformational change which could be accompanied by tilting of the second α-helix in the CHAP domain (α4 in Extended Data Fig. 3a) or by movement of the D-Ala C_α_ atom (Extended Data Fig. 3c). The oxygen atom in the D-Ala carboxyl is coordinated by hydrogen-bonds to Asn183 and Gln97. The two residues likely form the oxyanion hole which stabilizes the negative charge of the transition state forming during peptide hydrolysis (Fig. 3b).

The remainder of the stem peptide is coordinated by interactions with residues Asp91, Phe94, Phe99, Leu114, Gly116, Asp117, Ala119 and Phe159 (Fig. 3b). The *Sa* stem peptide differs at positions 2 and 3 from other gram-positive and gram-negative bacteria, with a D-iso-Gln instead of D-iso-Glu and L-Lys instead of meso-diaminopimelic acid (meso-Dpm). The CHAP specificity for the *Sa* stem peptide may be conferred by residues Leu114 and Gly116, which interact with the free amino group of D-iso-Gln through backbone carboxyl oxygen atoms and Phe94/Tyr159, which sandwich the carbons of the L-Lys side chain through hydrophobic interactions (Fig.3b, Extended Data Fig. 3d).

Comparison of the different CHAP variants and states shows that substrate binding induces a conformational change in the loop harboring residues Asp117 and Tyr118, which “flops” on the substrate, and Phe94/Phe99/Tyr159, which re-orient their hydrophobic sidechains to accommodate the substrate (Extended Data Fig. 3d).

Crystallization of full length LysM-CHAP variant L1-3 after incubation with Pep1a-CMK allowed us to additionally solve the structure of the full-length enzyme at medium resolution (2.8 Å). The structure shows how the LysM and CHAP domains align to form a continuous channel which may accommodate a complete peptidoglycan fragment consisting of a NAG-NAM chain and the stem peptide, as suggested by structural alignment of the LysM-glycan complex from NlpC^4^ with the LysM domain in the L1-3 structure (Fig. 3a). The interface formed by the two domains is mostly composed of amino acids 30-43 in the LysM domain and amino acids 90-93 and 111-117 in the CHAP domain. Key hydrogen bonds are formed between Lys13, Ser93 and Gly92 and between Asn41, Phe44 and Tyr117 (Extended Data Fig. 3e). Interestingly, the LysM domain contributes to the interaction with the stem peptide via backbone interactions with residue Gly14, Ile43 and Ile45 (Extended Data Fig. 3e), which may explain why it was only possible to determine full length structures of LysM-CHAP derivatives in the presence of substrate. This finding implicates that productive substrate binding of LysM-CHAP is a cooperative process which requires the LysM and CHAP domains and the substrate to come together simultaneously in the correct orientation.

Native peptidoglycan is further composed of the pentaglycine bridge that follows D-Ala (positions 5-9 in Extended Data Fig. 3b). The continuous cleft formed by the LysM and CHAP domains extends beyond the P1 site at Cys98 and may contribute to additional weak interactions with the substrate prior its cleavage. Since substrate-bound structures could only be determined with a covalently attached stem peptide analogue, it is likely that the interactions promoting the formation of the ternary LysM-CHAP-substrate complex are individually very weak and transient. We hypothesize that cooperative binding involving multiple weak interactions between LysM, CHAP and substrate allow the enzyme to bind to intact peptidoglycan with high affinity. However, once peptidoglycan is hydrolysed, the loss of some of these interactions causes the enzyme to rapidly dissociate from the product, allowing it to engage to an intact peptidoglycan cross-bridge and initiate a new hydrolysis cycle (see model in Ext. Fig. 3h). The reduced affinity to the product may further prevent product-inhibition of the enzyme once most of the peptidoglycan cross-links have been hydrolysed.

**Synergy with antibiotics in vitro and in vivo**

Interactions between lysins and antibiotics are gaining importance, especially considering potential clinical trials designs^5^. Several studies demonstrate synergies with different classes of antibiotics such as vancomycin or daptomycin^6-8^.

We demonstrated synergy with cell wall active antibiotics in several *in vitro* settings (Fig. 4a-d). Interestingly, while testing lysin/antibiotic combinations against different staphylococcal species (Extended Data Fig. 5a), we observed that co-treatment with vancomycin enabled eradication of *S. capitis*, which could not be eliminated by the lysin alone at up to 160 µg/mL. In contrast, dose-dependent antagonism between lysin and vancomycin was observed for *S. lugdunensis*, which may be related to peptidoglycan cross-bridge remodeling by its autolysin IsdP^9^. When treating *S. lugdunensis* with a combination of the cell-wall acting antibiotic cefazolin and lysin, synergy was observed. These findings emphasize the need for careful design of clinical trials and thorough testing of combinations in a pre-clinical setting.

Vancomycin was titrated in combination with a constant concentration of Fc-hel8-L1-3 or Fc-hel14-L1-6 (Extended Data Fig. 5b). Synergy was not only observed at the reported vancomycin plasma levels in humans (15-25 µg/mL), but across the vancomycin concentration series down to 2 µg/mL. These findings are particularly promising, as synergistic effects might be maintained over time-dependent fluctuating vancomycin exposure levels *in vivo*.

Other non-Fc containing lysin architectures such as CHAP-SH3 (exebacase), PepM23-SH3 (lysostaphin) or non Fc-fused LysM-CHAPs showed less or no synergy with vancomycin (Extended Data Fig. 5c). We postulate that the maximum (or close to maximum) possible peptidoglycan degradation may be reached by these lysins and thus there is no additional benefit of the addition of vancomycin.

An interesting avenue, which was not explored in this study, would be the assessment of bactericidal activity of lysin-antibiotic combinations on Vancomycin intermediate resistant Staphylococcus aureus (VISA) strains. Adding lysin to the treatment could potentially lead to a resensitization of these strains to vancomycin (e.g. by degradation of the thickened cell wall^11^ by lysin) and thus improve the therapeutic outcome.

*In vivo* efficacy studies were performed in mice with lysin dosing multiple times starting 1 hpi (Fig. 6a). Delayed treatment start at 13 hpi, as was performed in the combination studies (Fig. 6b), led to a decrease in efficacy for the lysin alone as well as for vancomycin. For RiboLysin administrations 1 hpi, treatment start is also somewhat delayed, as Cmax levels are reached after 7-8 hours (Fig. 6a), This could be an avenue to explore the postulated Fc-dependent mode of action where intra-macrophage killing by the lysin improves bacterial clearance.
